# Scribble controls social behaviors through the regulation of the ERK/Mnk1 pathway

**DOI:** 10.1101/2020.09.10.289397

**Authors:** Maïté M. Moreau, Susanna Pietropaolo, Jérôme Ezan, Benjamin J.A. Robert, Sylvain Miraux, Marlène Maître, Yoon Cho, Wim E. Crusio, Mireille Montcouquiol, Nathalie Sans

## Abstract

Social behavior is a basic domain affected in several neurodevelopmental disorders. Indeed, deficits in social interest, interactions and recognition represent core symptoms of Autism Spectrum Disorder but are also found associated with a heterogeneous set of neuropsychiatric and rare disorders. The *SCRIB* gene that codes for the polarity protein SCRIBBLE has been identified as a risk gene for spina bifida, the most common type of open neural tube defect, found at high frequencies in autistic patients, as other congenital anomalies, while the deletions/mutations of the 8q24.3 region encompassing *SCRIB* genes is associated with multisyndromic and rare disorders. Nonetheless, the potential link between *SCRIB* and ASD-relevant social phenotypes has not been investigated yet. Hence, we performed an extensive behavioral characterization of the *circletail* line that carries a mutated version of *Scrib*. Scrib^crc/+^ mice displayed reduced social interest, lack of preference for social novelty and social reward, and reduced social habituation while other behavioral domains were unaltered. Social deficits were associated with reduced hippocampal volume, upregulation of ERK phosphorylation in specific hippocampal regions, together with increased c-Fos activity in the same brain areas. Importantly, the social alterations were rescued by both direct and indirect pERK inhibition. These results support a specific link between polarity genes, social behaviors and hippocampal functionality, thus suggesting a role for *SCRIB* in the etiopathology of neurodevelopmental disorders. Furthermore, our data demonstrate the crucial role of the MAPK/ERK signaling pathway, in underlying the social deficits induced by *SCRIB* mutation, thus supporting its relevance as a therapeutic target.

## Introduction

Social approaches, interactions and relationships are the basis of our normal and modern life (Adolphs, 2009). These social relationships rely on social memory and are built on individual abilities to differentiate other person, to remember them in order to have the appropriate behavioral responses based upon previous encounters. Dealing with these social situations can be considered as problematic for some individuals. Given the importance of social interactions in our everyday life, it is crucial to understand their bases at molecular, neural and network levels to develop new pharmacotherapeutics that can ease dysfunctional social interactions. To some extent, these social dysfunctions become characteristic symptoms of various psychiatric illnesses, including several neurodevelopmental disorders (NDDs) and genetic disorders such as autism, but are also found in mood and anxiety disorders that probably stem their origin in prenatal life during fetal development (Courchesne et al., 2019).

*SCRIB* gene (*SCRIBBLE, SCRB1*; OMIM# 607733), located on chromosome 8, encodes for a protein named Scribble (Scrib or Scrib1) which is key in many developmental process including cell proliferation, migration, and polarity (Bonello and Peifer, 2019). Scrib is also known for its involvement in planar cell polarity signaling that ultimately regulates embryonic and postnatal development (Montcouquiol et al., 2003, 2006a; Moreau et al., 2010; Murdoch et al., 2003). Mutations in *SCRIB* are described in patients with spina bifida (Lei et al., 2013; Robinson et al., 2012; Wang et al., 2019), one of the most common forms of neural tube defect, found at high frequency in patients of Autism Spectrum disorder (ASD) (Dawson et al., 2009; Lauritsen et al., 2002; Timonen-Soivio et al., 2015). Indeed, accumulating evidence suggests a role of *SCRIB*, together with other polarity genes, in neurodevelopmental and autistic disorders (for review, see (Sans et al., 2016)). Mutations 8q24.3 in the locus encompassing the *SCRIB* gene have been linked to the rare Verheij syndrome (VRJS, OMIM #615583) characterized by intellectual disability (Dauber et al., 2013) and to ASD as well (Hu et al., 2015; Iossifov et al., 2012; Neale et al., 2012). Moreover, Scrib is also known to play a role in the modulation of the MAPK signaling and to interact directly with ERK (Nagasaka et al., 2010). Chromosomal deletions encompassing ERK-encoding *MAPK* genes are commonly associated with neurodevelopmental disorders (NDDs) and ASDs (Campbell et al., 2008; Fernandez et al., 2010; Saitta et al., 2004). Furthermore, abnormalities in ERK functionality have been described in several mouse models of NDDs and ASD (Bagni and Zukin, 2019; Courchesne et al., 2019; Faridar et al., 2014; Vithayathil et al., 2018). Mouse studies have also shown a major role of Scrib in modulating dendritic arborization and connectivity, post-endocytic NMDA receptor trafficking and hippocampal synaptic plasticity (Moreau et al., 2010; Hilal et al., 2017), i.e. processes that are known to play a key role in the etiopathology of in NDDs and ASD (Bagni and Zukin, 2019; Cheng et al., 2017; Lin et al., 2016). The potential link between *SCRIB*, brain development and function and underlying ASD-like behaviors are poorly understood. Also, understanding of the molecular systems involved in the molecular pathology of *SCRIB* -associated neurodevelopmental disorders including autism form the bases of ongoing work to build autism biomarkers (Diaz-Beltran et al., 2016).

To fully understand the physiological and pathophysiological mechanisms regulated by Scrib, we conducted an extensive neurobehavioral characterization of heterozygous *circletail* mutant mice (Scrib1^*crc/+*^), carrying a mutation in *Scrib* to investigate a potential link between this polarity gene and ASD-like behaviors. Our focus was on social behaviors (social interest, preference for social novelty and social reward in the three-compartment test, direct social interaction with assessment of ultrasonic communication, social habituation), as they represent core symptoms of ASD that are also common to multiple NDDs. Non-social behaviors were assessed using tests for locomotion (open field), emotionality (Elevated plus-maze, dark-light box and neophobia), repetitive behaviors (marble, grooming), olfactory discrimination/habituation, sensory-motor responsiveness (acoustic startle and its prepulse inhibition) cognitive abilities (based on novelty preference, i.e., object recognition and spontaneous alternation in the Y maze). This variety of tests allowed us to evaluate whether the potential impact of Scrib mutation on social behaviors could (i) be confounded by the interference of other behavioral alterations (e.g., changes in general activity, emotionality, olfaction), (ii) be specific for this domain or instead affects general novelty preference and detection (e.g., object recognition, spontaneous alternation, olfactory habituation). We then investigated the potential brain mechanisms underlying the behavioral effects of Scrib mutation using MRI and c-Fos expression approaches, as well as assessing the functionality of the MAPK/ERK pathways. Finally, we investigated whether the behavioral deficits of Scrib^*crc/+*^ mice could be specifically rescued by inhibiting the MAPK/ERK pathway through systemic administration of MEK and Mnk1 inhibitors. Mnk1 kinase is a known downstream target of ERK signaling and a recent study suggested it could be a part of a molecular signature for ASD (Lim et al., 2013; Rosina et al., 2019). Importantly, we were able to specifically rescue these behavioral deficits by systemic administration of MEK and Mnk1 inhibitors of the MAPK/ERK pathway in adult mice. This supports the hypothesis that the MAPK/ERK pathway is hyperactive in the Scrib^*crc/+*^ mice, and this underlies the sociability deficits. Altogether, our results suggest that social behaviors are controlled by a Scrib regulation of the ERK-Mnk1 overlapping mechanisms of phosphorylation.

## Results

### Scrib^crc/+^ mice have deficits in social interest and lack of preference for social novelty and social reward

To determine whether the *Scrib*^*crc/+*^ mouse has social deficit, we first tested the mice in the social interest and preference for social novelty paradigm using the three-compartment test (Figure 1A). During habituation, the time spent in the right/left chamber during habituation was not different between the WT and *Scrib*^*crc/+*^ (no genotype x chamber interaction: *F*_1,34_ < 1, n.s; Figure S1A). In the social interest test (Figure 1C), WT mice had a significantly higher preference for the stranger male mouse (S1, social stimulus) than *Scrib*^*crc/+*^ animals (genotype x stimulus interaction; *F*_1,34_ = 21.34, *p* < 0.0001). Accordingly, WT spent more time spent around the small wire cage containing the stranger (S1) than the non-social novel stimulus (O) (*t*_16_ = 6.57, *p* < 0.0001; Figure 1C), whereas *Scrib*^*crc/+*^ mice showed no preference for the stranger male mouse (*t*_18_ = 0.22; ns; Figure 1C). At the end of the social interest test, each mouse was tested for social novelty preference when a novel social stimulus (S2) replacing the non-social one (O) (Figure 1D). WT mice showed a clear preference for the compartment containing the novel (S2) rather than the familiar social stimulus (S1), while this preference was not observed in *Scrib*^*crc/+*^ mice, which spent approximately the same amount of time in the two compartments (genotype x stimulus interaction: *F*_1,34_ = 4.88, *p <* 0.05; S1 vs. S2, Bonferroni corrected *t-*test: WT: *t*_16_ = 3.84, *p <* 0.001; *Scrib*^*crc/+*^: *t*_18_ = 0.84, n.s; Figure 1D). For all trials, the number of entries in both stimulus chambers was similar for both genotypes (Figure S1B-D), suggesting no difference in general exploration levels. All together these results show that *Scrib*^*crc/+*^ mice have a deficit in the natural preference for social interaction and in social novelty recognition.

**Figure 1.**
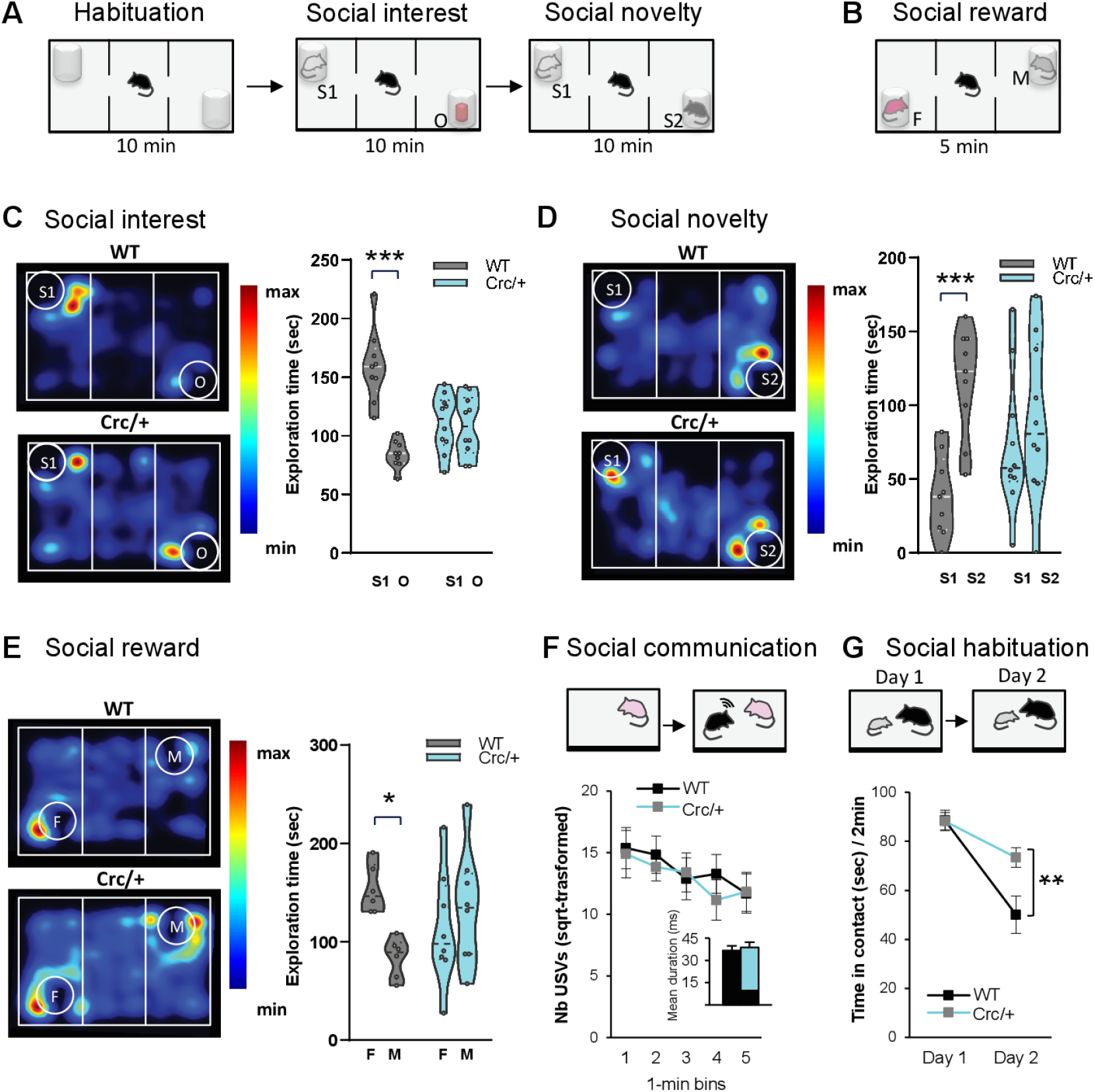
*Scrib1*^*crc/+*^ mutant mice display selective deficits of social interactions. **A-B** Experimental design protocol of the three-chambered choice test. **A, C** Time spent sniffing with stranger male mice (S1, social stimulus) versus the object (O, non-social stimulus) during the social interest test. Here, control show a preference for the stranger mice while *Scrib1*^*crc/+*^ mice show no preference between the stranger mice and the object. **A, D** Time spent sniffing with novel male mice (S2) versus familiar male (S1) mice during the social novelty test. Control group show a high preference for the novel stranger mice but *Scrib1*^*crc/+*^ mice show no preference between the novel stranger mice and the familiar male mice. **B, E** Time spent sniffing a male (M) versus a female (F) mouse during the social reward test. Controls show a preference for the female mice during the first 5 min of the test while *Scrib1*^*crc/+*^ mice did not. **F** Experimental design protocol of the direct social interaction with female during 5 min. Mean of duration and number of ultrasound vocalizations during a 5 min test. **G** Experimental design protocol of the direct social interaction/habituation with a juvenile. Mean investigation duration with a juvenile during initial social recognition (day 1) and social habituation (day 2). For c, d, and e data is presented as median with 25^th^ and 75^th^ percentile and single data points are shown as dot. For f and g data is presented as mean ±SEM. The shaded area represents the probability distribution of the variable. \**p* ≤ 0.05; \*\**p* ≤ 0.01 and \*\*\**p* ≤ 0.001

We then assessed the preference for a more attractive, rewarding stimulus, i.e., an adult female (F), in the three-compartment apparatus compared to a stranger male mouse stimulus (M) (Figure 1B). During the test, WT mice showed a preference for the female stimulus (F), which was absent in mutants (genotype x stimulus interaction: *F*_1,24_ = 6.447, *p <* 0.018; Bonferroni corrected *t-*test: WT: *t*_24_ = 2.58, *p <* 0.05; *Scrib*^*crc/+*^: *t*_24_ = 0.89, n.s; Figure 1E). These results show that *Scrib*^*crc/+*^ display lack of preference for social reward.

### Scrib^crc/+^ mice display deficits in social habituation with a juvenile but normal social interaction and communication with adults

Next, we used a direct social interaction test to assess ultrasonic vocalizations during the direct social behavior. The time spent in social investigation was similar between *Scrib*^*crc/+*^ mice and their WT littermates indicating that *Scrib*^*crc/+*^ mice have no major defect in direct social interaction with an adult female (*t*_*24*_ = 0.16, n.s; Figure S1E). No difference in the ultrasonic vocalizations emitted during the social interaction session was observed, in terms of both number and mean duration (no genotype effect: *F*_1,22_ < 1, n.s.; Figure 1F), indicating that *Scrib*^*crc/+*^ mice have no communication defects with an adult female.

Next, we tested the mutant mice in a social habituation paradigm (Figure 1G) that provides a measure of hippocampal- and amygdala-dependent social memory (Kogan et al., 2000). As expected, WT and *Scrib*^*crc/+*^ mice showed a characteristic social habituation, i.e. decrease in duration of social investigation with a juvenile male during the two repeated presentations of the same stimulus mouse (Day effect: *F*_1,48_ = 5.63, *p* < 0.05). In contrast, *Scrib*^*crc/+*^ mice presented a slower habituation compared to WT (genotype x day interaction: *F*_1,48_ = 5.97, *p* < 0.05; Bonferroni corrected *t*-test: Day 2 comparison Test, WT vs *Scrib1*^*crc/+*^; *t*_24_ = 3.40, *p* < 0.01; Figure 1G). These data indicate that *Scrib*^*crc/+*^ mice have normal direct social interaction with a juvenile mouse, but have a deficit in social habituation.

### Scrib^crc/+^ mice do not have repetitive behavior or sensorimotor gating defect

To determine whether the social behavior deficits were accompanied by other type of difficulties, we first investigated potential repetitive behaviors. The *Scrib*^*crc/+*^ mice showed normal rearing (WT vs. *Scrib*^*crc/+*^, *t*-test, *t*_*23*_ = 0.011, n.s; Table S1) and self-grooming in their home cages, similar to control mice (WT vs. *Scrib*^*crc/+*^, *t*-test, *t*_*23*_ = 0.68, n.s; Table S1). Also, no differences between groups were found when comparing the number of repetitive entries in the same arm of the Y-maze test (percentage correct alternations; WT vs. *Scrib*^*crc/+*^, *t*-test, *t*_*23*_ = 0.37, n.s; Table S1), and the number of repetitive digging behavior in the marble-burying test (WT vs. *Scrib*^*crc/+*^, *t*-test, *t*_*23*_ = 0.44, n.s; Table S1). Using the startle response and prepulse inhibition (PPI), we found that *Scrib*^*crc/+*^ had normal sensory-motor gating (pulse effect for WT and *Scrib*^*crc/+*^respectively, *F*_2,4_ = 269.98 and 10.05, n.s; Table S2). Altogether, these results did not reveal problem of stereotyped behaviors and sensory-motor integration in *Scrib*^*crc/+*^ mice.

### The social deficits in Scrib^crc/+^ mice are not a consequence of anxiety, alterations in locomotion, or neophobia

The social behavior deficits found in the *Scrib*^*crc/+*^ mice could be the result of increased anxiety, altered locomotion, neophobia, or impairment in general novelty recognition which are all crucial for social interest and novelty. We did not detect a deficit in any of these behaviors when we submitted the *Scrib*^*crc/+*^ mice to a battery of test (Table S1). We also used a familiar open field environment to ascertain the intact abilities of WT and mutant mice to explore novel objects (Figure S2A-C) as described previously (Dulawa et al., 1999). In the open field, both WT and *Scrib*^crc/+^ mice completed a high number of entries and spent more time in the center after introduction of the object (effect of object: F_1,44_ = 52.23, p < 0.0001*** for entries; F_1,44_ = 46.96, p < 0.0001*** for time; no genotype effect: F_1,44_ = .75, n.s for time; F_1,44_ = 0.06, n.s for entries; Figure S2A), confirming that the novel object elicited curiosity.

These data suggest that *Scrib*^*crc/+*^ mice have normal behavioral responses to novelty in the object exploration test. Next, we showed that both groups of mice spent a comparable amount of time exploring the environment with two identical objects (no object effect: *F*_1,24_ < 1, n.s; no genotype effect: *F*_1,24_ < 1, n.s; Figure S2B). When one object was replaced by a new object, both groups exhibited a significant preference for the novel object (Object effect: *F*_1,24_ = 12.84, *p* < 0.01**; no genotype effect: *F*_1,24_ < 1, n.s; no genotype x object interaction: *F*_1,24_ < 1, n.s; Figure S2C). These data indicate that *Scrib*^*crc/+*^ and WT mice have a normal object-recognition memory and do not display neophobia.

### Scrib^crc/+^ mice have normal olfactory discrimination

Since social behavior in mice is strongly dependent on olfaction (Farbman, 1994; Guillot and Chapouthier, 1996), we evaluated whether *Scrib*^*crc/+*^ mice have olfactory deficits. We tested both WT and *Scrib*^*crc/+*^ mice in an olfactory-guided foraging task (Moy et al., 2004). The time required to locate the food buried under the sawdust was similar for both genotypes without food deprivation conditions (WT vs. *Scrib*^*crc/+*^, *t*-test, *t*_*26*_ = 0.40, n.s; Figure S3A) or following food deprivation conditions (data not shown). Habituation and dishabituation (Figure S3B) were observed for social and non-social odor cues with no statistical difference between WT and *Scrib*^*crc/+*^ mice (no genotype effect: *F*_1,187_ = 1.87, n.s). Both genotypes displayed similar levels of habituation indicated by a decrease in the time spent sniffing the odorant stimulus following its repeated presentation and comparable levels of dishabituation, indicated by increased time sniffing a novel odorant stimulus (odor effect: *F*_11,187_ = 22.54, *p* < *0*.*0001*; no odor x genotype interaction: *F*_11,187_ = 1.30, n.s; Figure S3B). These data indicate that *Scrib*^*crc/+*^ mice do not have olfactory deficits. Altogether, our data rule out abnormal motor function, general novelty recognition problems, or sensory alterations as the cause of the reported social deficits in the *Scrib*^*crc/+*^ mutant mice.

### c-Fos levels are increased in the DG and CA3 region of the hippocampus of Scrib^crc/+^ mice after social exposure

To specifically map brain regions activated or not during the social interest test and potentially affected by Scrib loss, we first performed c-Fos and *Zif268* neuroimaging (Figure 2). Both WT and *Scrib*^*crc/+*^ mice were subjected to the social interest test in the three-compartment chamber and left undisturbed in their home cages for 1 h before perfusion (Social condition). In our control control group, both WT and *Scrib*^*crc/+*^ mice were placed in the empty three-compartment chamber for the same time and then returned to their home cages until tissue collection (Control condition). In WT mice, after the social interest test, we observed an increase in the density of c-Fos-positive cells compared to the control condition in restricted areas of the basal forebrain including hippocampus areas (DG, CA1, CA3), enthorinal cortex (EntCx), medial nucleus of the amygdala (MeA), motor (MCx) and piriform cortex (PirCx), striatum (ST) as well as in the granular cells of the olfactory bulb (MOB Gcells) (Figure S4A-B). In *Scrib*^*crc/+*^ mice, this pattern was modified. After the social interest test, we found that the same regions were activated as in the WT group but also the density of c-Fos-positive cells was significantly increased in the Sensorimotor cortex (S1Cx) and in the cortical nucleus of the amygdala (CoA). On the other hand, the dorso lateral part of the striatum (STDL) was not activated in the *Scrib*^*crc/+*^ mutant after social interest test compared to the *Scrib*^*crc/+*^ mice control group (Fig 2B and Figure S4B). Furthermore, in both control and social interest condition this pattern was modified specifically by the mutation. We found that the number of c-Fos -positive cells measured were significantly increased in the olfactory bulb (AOB/MOB) and decreased in the piriform cortex (PirCx) in the *Scrib*^*crc/+*^ mutant compared to WT (Fig 2B and Figure S4C-D). These observations show that c-Fos activity in these structures is modulated by Scrib expression levels but are independent of the social behavior task. However, after social interest test a more robust increase of number of c-Fos - positive cells expression was observed in the amygdala (CoA, ≈+56%), the DG (≈+12%) and CA3 (≈+55%) region of the hippocampus and the Cortex (S1, ≈+48,7%) in *Scrib*^*crc/+*^ compared to WT mice but not in the CA1 or the medial nucleus of the Amygdala (MeA) (Fig 2D and Figure S4D). This up-regulation of c-Fos levels in the CA3 and DG of the *Scrib*^*crc/+*^ mice was the direct consequence of the social interest exposure, because the expression levels of c-Fos in the control animals were comparable between the WT and the *Scrib*^*crc/+*^ mice in these two regions (Figure 2D). In the CA3 and DG of the hippocampus, we found a similar up-regulation of *Zif268*-positive cells (Table S3). These results show that the DG and CA3 regions of the hippocampus are the main activated ones during the social interest test in the *Scrib*^*crc/+*^ mutant mice.

**Figure. 2.**
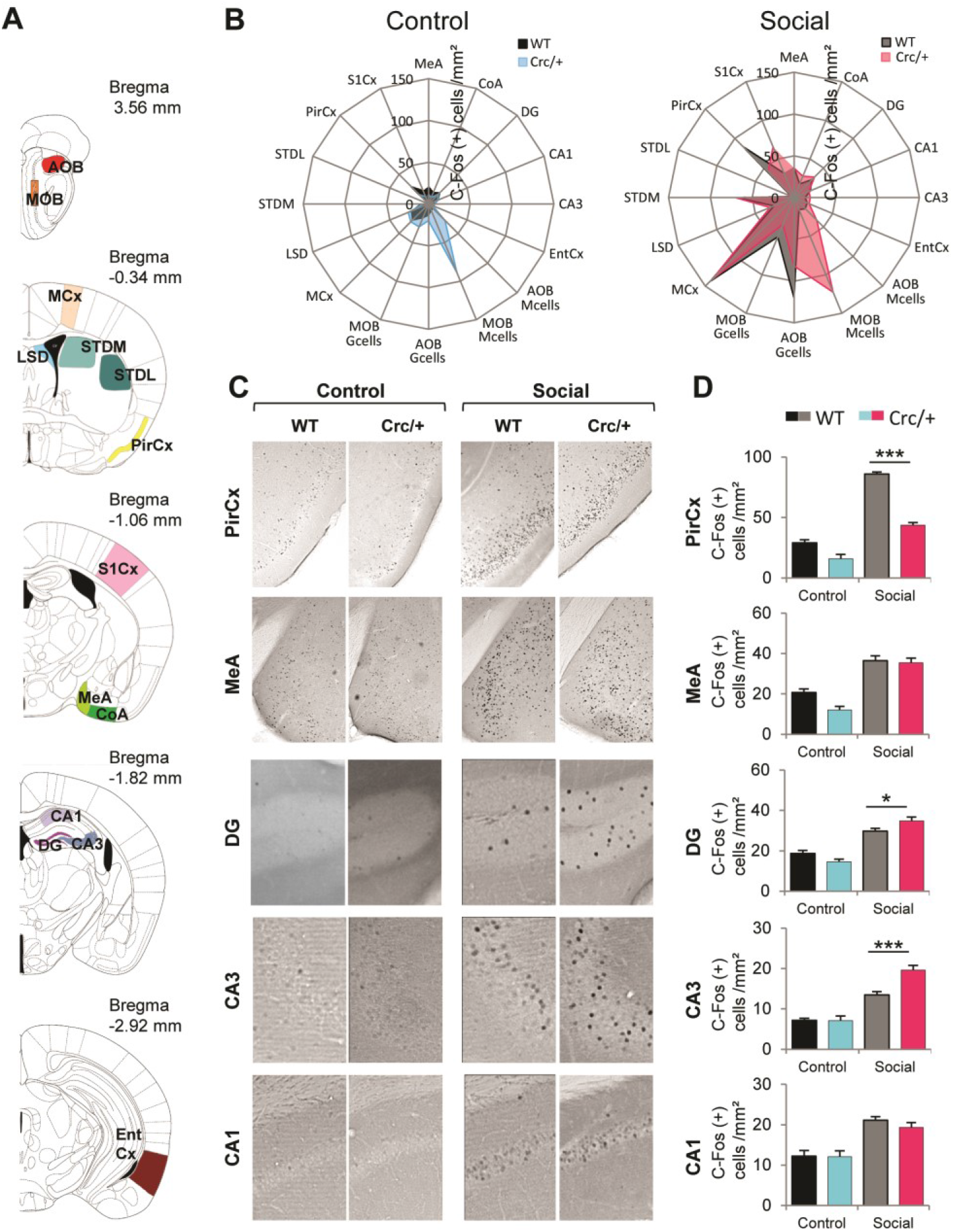
Specific patterns of neural activation after social interest test in *Scrib1*^*crc/+*^ mice. **A** Coronal section of neuroanatomical areas analyzed for c-Fos immunoreactivity after a social interest test adapted from Paxinos and Franklin (1997). Accessory olfactory bulb (AOB) and the main olfactory bulb (MOB) granular (Gcells) and mitral cell subdivisions (Mcells); the motor cortex (MCx); the dorsal portion of the lateral septum (LSD); the dorso medial (STDM) and the dorso lateral (STDL) part of the striatum; the piriform cortex (PirCx); the somatosensoriel cortex (S1Cx); the medial nucleus (MeA) and the cortical nucleus of the amygdala (CoA); the CA1 (CA1) and the CA3 (CA3) subregion of the hippocampus; the dentate gyrus (DG) and the lateral entorhinal cortex (EntCx). **B** Radar chart recapitulate the activate brain region as measured by change in c-Fos immunoractive cells in WT and *Scrib1*^*crc/+*^ mice in the control and social condition. **C** Representative microphotographs showing c-Fos-positive cells (dark dots) in the PirCx, MeA, DG, CA3 and CA1 of WT mice or *Scrib1*^*crc/+*^ mice that were killed 1 hour after the control/social test. **D** Number of c-Fos-positive cells per mm^2^ (mean ± SEM) was calculated for each genotype and exposure condition. All data are presented as mean ±SEM. \**p* ≤ 0.05 and \*\*\**p* ≤ 0.001.

### Reduced structural integrity of the hippocampus hippocampus of Scrib^crc/+^ mice

Next, we investigated whether or not the behavioral alteration associated with c-Fox expression was accompanied by a change in size/volume of brain regions. Magnetic Resonance Imaging (MRI) analysis showed no major changes in the global volume of the brain (Figure 3), confirming a previous study (Moreau et al., 2010). However, MRI revealed an increase in the volume of the olfactory bulbs and a decrease in the volume of the hippocampus (Figure 3A,B) that was confirmed using 3D reconstruction analysis on brain sections of *Scrib*^*crc/+*^ mutant mice (Figure 3C,D). Together, these findings point to an alteration in the structure and /or connectivity between the OB maybe to compensate the defects the hippocampus in the mutant mice.

**Figure 3.**
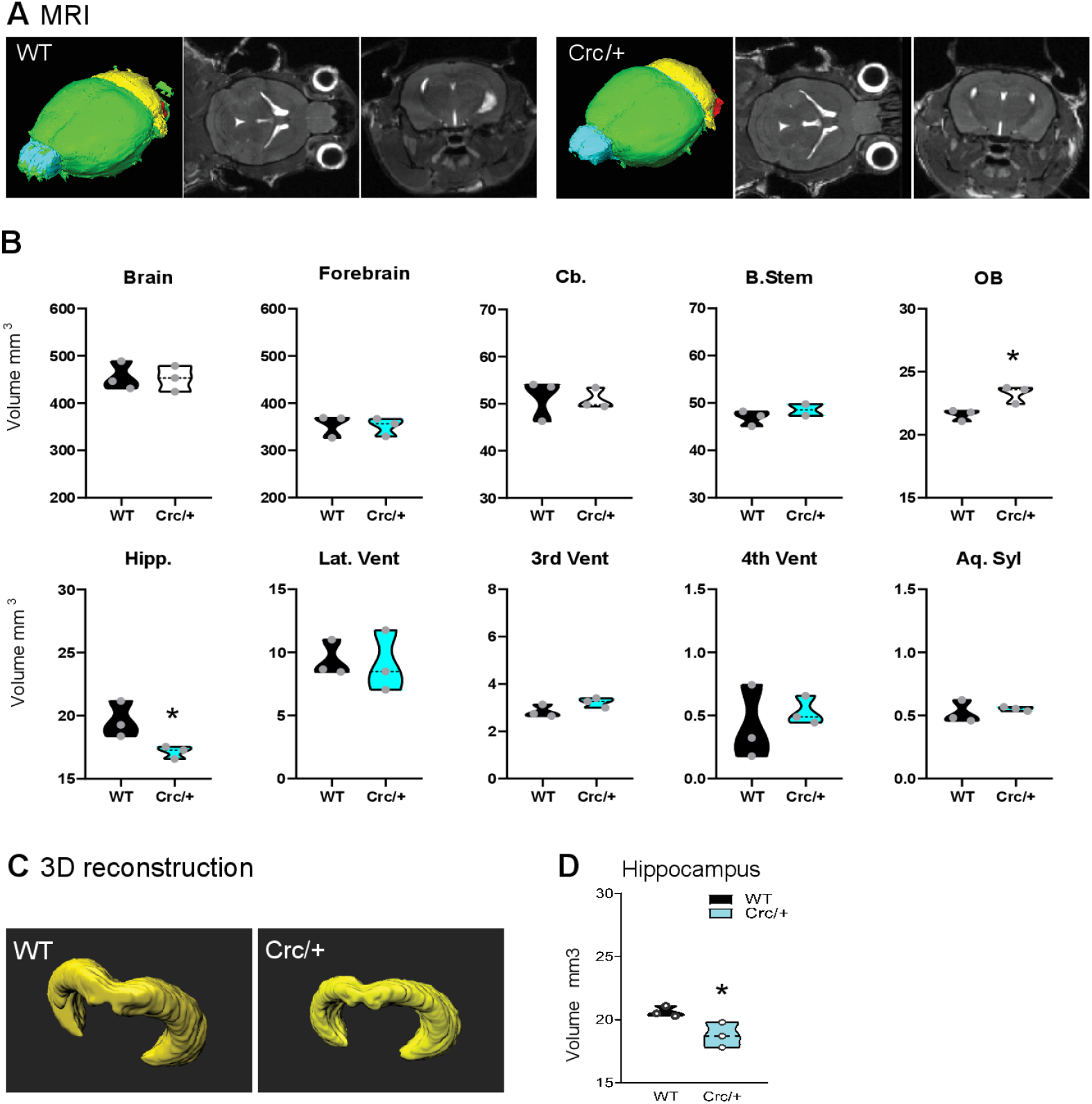
Scrib1 mutation decreases hippocampus volume. **A** MRI 3D reconstruction (left); sagittal (middle) and coronal (right) maps of a WT and *Scrib1*^*crc/+*^ mice brain in vivo. **B** MRI volume quantification show a specific reduction of the hippocampus volume and increased olfactory bulb volume in *Scrib1*^*crc/+*^ mice brain. **C** Imaris 3D reconstruction of a WT and *Scrib1*^*crc/+*^ mice hippocampus. **D** Volume quantification show reduction of hippocampal formation in the *Scrib1*^*crc/+*^ mice. All data are presented as median with 25th and 75th percentile and single data points are shown as dot. The shaded area represents the probability distribution of the variable. \**p* ≤ 0.05.

### Reduction of Scrib levels leads to significant increase in ERK phosphorylation in the hippocampus of Scrib^crc/+^ mice after social exposure

Scrib can inhibit the activation of extracellular signal regulated kinase (ERK) pathways in various systems (Bonello and Peifer, 2019; Jarjour et al., 2015; Nagasaka et al., 2010) and ERK and the mitogen activated protein kinase (MAPK) signaling pathway are linked to neurodevelopmental disorders and more specifically ASD (Bagni and Zukin, 2019; Courchesne et al., 2019; Faridar et al., 2014; Vithayathil et al., 2018). Thus, we assessed the level of activated phosphorylated ERK in the three subfields of the hippocampus of *Scrib*^*crc/+*^ mice in response to the social interest test. Our data show that ERK1 and ERK2 were activated in the three regions of the hippocampus of both mice the mice which performed the test (both *Scrib*^*crc/+*^ and WT) compared to the unexposed group (Figure 4A-C). Two-way ANOVA indicated a main effect of the social test exposure on the p-ERK1 levels on both genotype of each subfield (Social test effect, CA1: *F*_1,12_ = 16.24, *p* < 0.01; CA3: *F*_1,12_ = 12.47, *p* < 0.01; DG: *F*_1,12_ = 6.09, *p* < 0.05; Figure 4B). Exposure to the test also significantly increased ERK2 phosphorylation levels in both genotype in CA3 (Social test effect, CA1: *F*_1,16_ = 0.11, *n*.*s*; CA3: *F*_1,15_ = 10.65, *p* < 0.01; DG: *F*_1,16_ = 4.131, *n*.*s*; Figure 4C). On the other hand, the reduction of Scrib levels in the *Scrib*^*crc/+*^ mice leads to a hyper-activation of ERK during social exposure in the DG and the CA3, but not in the CA1 region, compared to WT mice performing the same test (genotype x exposure interaction: pERK1-CA3: *F*_1,12_ = 6.41, *p* < 0.05; pERK2-DG: *F*_1,16_ = 8.96, *p* < 0.01; pERK2-CA3: *F*_1,15_ = 9.88, *p* < 0.01). In conclusion, ERK phosphorylation was significantly higher in the *Scrib*^*crc/+*^ mice hippocampus compared to WT after the social interest test (Figure 4B and Figure 4C).

**Figure 4.**
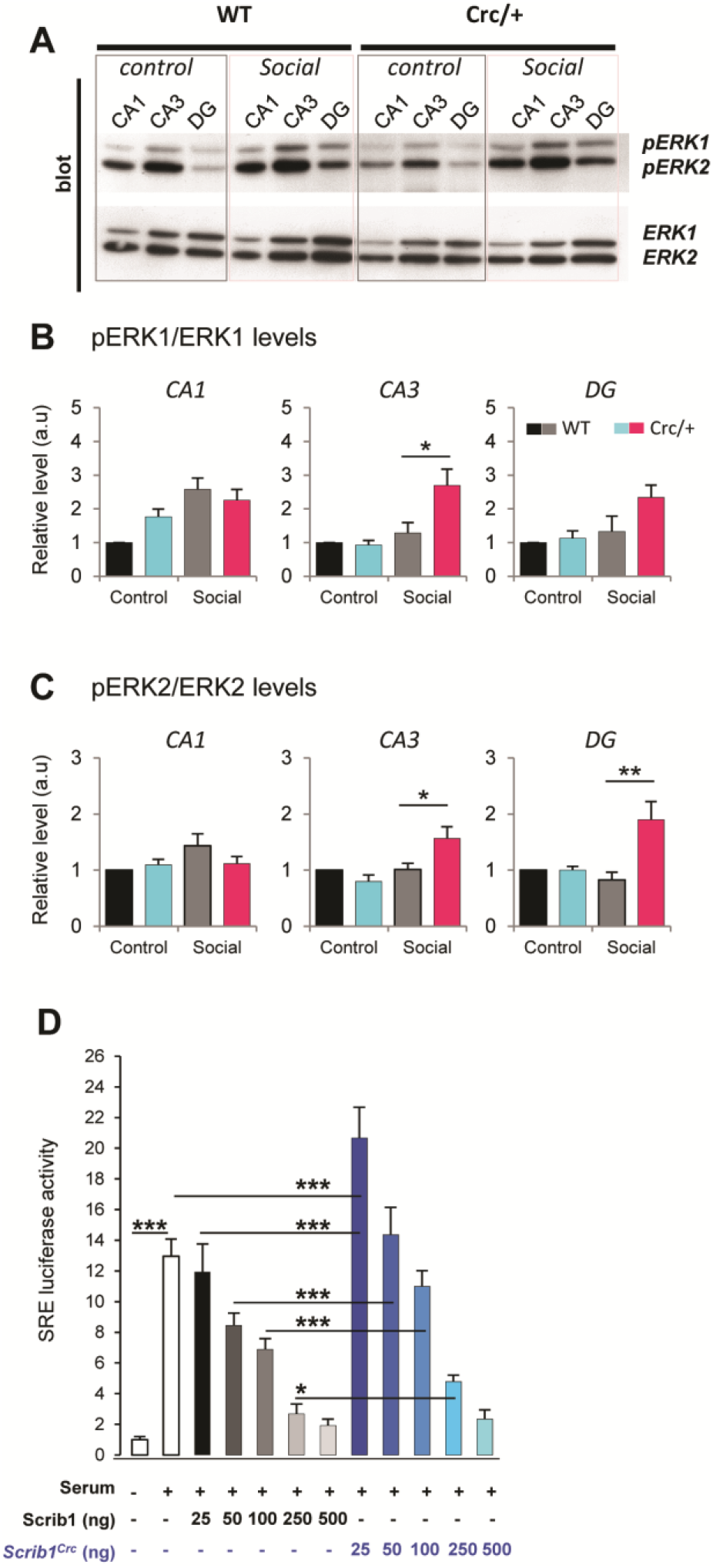
Scrib1 mutation modulates ERK proteins levels in the hippocampus and has a biphasic effect on the ERK signaling pathway. **A** CA1, CA3 and DG lazer capture homogenates from control and social WT and *Scrib1*^*crc/+*^. Mice are analyzed simultaneously for phospho-ERK1/2 (pERK1/2) and ERK1/2 by quantitative Western blotting. **B** Levels of phospho-ERK1 and **C** phospho-ERK2 in CA1, CA3 and DG. In control mice and *Scrib1*^*crc/+*^ mice, ERK1 and ERK2 phosphorylation level is significantly more elevated after social interest in CA3 and DG of *Scrib1*^*crc/+*^ mice. **D** Luciferase reporter assay data from HEK293 cells transfected with the SRE-LUC reporter vector along with increasing amount of the Scrib1 or *Scrib1*^*crc*^ expression plasmid as indicated. Twenty-four hrs after transfection, cells were lysed after being serum-starved for 2hrs then activated for 6hrs with 20% FBS as indicated. Activation threshold of the serum-induced ERK pathway is indicated by the horizontal dashed line. Scrib1 represses the ERK-dependent transcriptional activation of the SRE-Luc promoter in a dose-dependent manner. *Scrib1*^*crc*^ displayed a biphasic dose-response curve with an increased activation of the SRE-LUC activity effect at 25ng, whereas a significant inhibitory effect was observed at 100ng and higher doses. All data are presented as mean ±SEM. \**p* ≤ 0.05 and \*\*\**p* ≤ 0.001.

### Scrib^crc^ form is crucial for ERK pathway activation

To confirm the direct link between Scrib levels and ERK activation, we used a classical ERK signaling pathway *in vitro* reporter assay using SRE-LUC as a reporter gene. As expected and illustrated Figure 4D, increased levels of full length Scrib reduced the serum-induced SRE-LUC reporter gene expression in a dose-dependent manner, supporting the inhibitory role of Scrib on ERK signaling (ANOVA, *F*_*11*.*132*_ = 217.4; *p* < 0.0001). The C-ter truncated form of Scrib (the one expected from Scrib^*crc*^ mutation) exhibited a biphasic response. Lower doses significantly increased the luciferase expression when compared to the same concentration of full length Scrib (Scrib *vs*. Scrib1^crc^; Bonferroni comparison *t-*test: 25 ng: *t*_*16*_ = 13.68, *p* < 0.001; 50 ng: *t*_*16*_ = 9.248, *p* < 0.001; 100 ng: *t*_*34*_ = 8.57, *p* < 0.001; 250 ng: *t*_*16*_ = 3.31, *p* < .05), while higher doses were inhibitory, similar to full length Scrib, although not to the same extent (for illustration, compare Scrib and Scrib^*crc*^ at 500 ng/ml; *t*_*16*_ = 3.31, n.s). These results are consistent with an inhibitory role of Scrib on the ERK pathway, an inhibition that is reduced in Scrib^*crc*^ mutants due most probably to the absence of one ERK binding site in the truncated protein. The mechanisms leading to the potentiating effect observed for lower levels of Scrib^*crc*^ are unclear at the moment, but it could be the result of a dominant-negative effect of Scrib^*crc*^ on full length Scrib. This dual effect is consistent with ERK hyperphosphorylation in the *Scrib*^*crc/+*^ mice, and could explain part of the complex phenotype in these mice.

### Specific inhibition of ERK rescues social deficits in Scrib^crc/+^ mice

The lack of social interest in *Scrib*^*crc/+*^ mutant mice is associated with abnormally high levels of activated ERK levels in the hippocampus. If true, we should be able to rescue that phenotype by inhibiting ERK over-activation in the hippocampus via the administration of an inhibitor of the mitogen-activated protein kinase (MEK) SL327, as done previously (Fernandez et al., 2008).

WT and mutants were assessed in the three-compartment test one hour after vehicle or 30 mg/kg SL327 treatment (Figure 5A). Each treatment group was compared with its own saline control. *Scrib*^*crc/+*^ treated with SL327 showed normal levels of social interest comparable to their treated WT littermates (genotype x stimulus x SL327; *F*_*1,60*_ = 10,36; *p* < 0.01**; O vs S1; Bonferonni’s multiple comparaisons test: WT-SL327, *t*_*7*_ = 7.87, *p* < 0.0001***; *Scrib*^*crc/+*^*-SL327, t*_*7*_ = 6.35, *p* < 0.0001***; Figure 5D). The effect is comparable to the WT in saline condition (WT-Saline, *t*_*11*_ = 8.80, *p* < 0.0001***; Figure 5D) and different from the *Scrib*^*crc/+*^ in the saline condition (*Scrib*^*crc/+*^*-saline, t*_*9*_ = 1.17, *n*.*s*; Figure 5D). The same rescued effects were observed on the preference for social novelty (genotype x stimulus x SL327 effect; *F* _*1,64*_ = 25.81; *p* < 0.0001***; S1 vs S2; Bonferonni’s multiple comparaisons test: WT-SL327, *t*_*8*_ = 4.20, *p* < 0.001***; *Scrib*^*crc/+*^*-*SL327, *t*_*8*_ = 6.90, *p* < 0.001***; Figure 5E).

**Figure 5.**
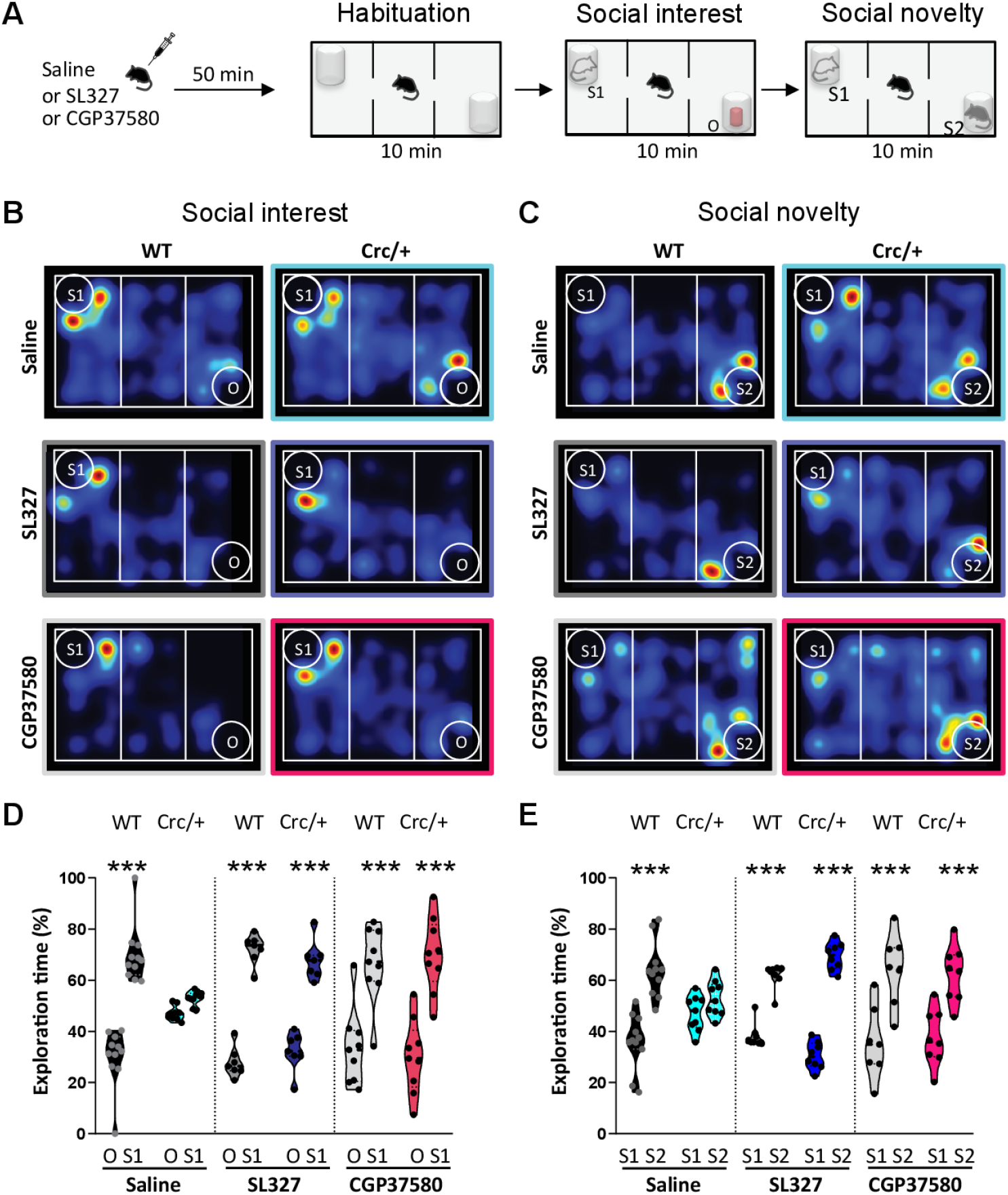
The inhibition of ERK in the *Scrib1*^*crc/+*^ mice reverses the autistic-like social deficits. **A** Experimental design protocol of the three-chambered choice test. **B, D** Time spent sniffing with stranger male mice (S1) versus object (O) during the social interest test after vehicle, SL327 or CGP37580 treatment. **C, E** Time spent sniffing with novel male mice (S2) versus familiar male mice (S1) during the social novelty test after vehicle, SL327 or CGP37580 treatment. Here, Scrib1 mutation causes a deficit in both social interest and social novelty test that is rescue by inhibition of ERK signaling. All data are presented as median with 25^th^ and 75^th^ percentile and single data points are shown as dot. The shaded area represents the probability distribution of the variable. \*\*\**p* ≤ 0.001.

### Dowstream of ERK signaling, the specific inhibition of Mnk1 in Scrib^crc/+^ mice also rescues social deficits

Mnk1 kinase is a known downstream target of ERK signaling and a recent study suggested it could be a part of a molecular signature for ASD (Bagni and Zukin, 2019; Lim et al., 2013). To determine whether the inactivation of Mnk1 could rescue the social deficits in the *Scrib*^*crc/+*^mice, we used CGP57380, a cell-permeable pyrazolo-pyrimidine compound that acts as a selective inhibitor of Mnk1, but has no inhibitory activity against ERK1/2 (Figure 5A). We found that Mnk1 inhibition improved social behavior in the *Scrib*^*crc/+*^ mice (genotype x stimulus x CGP57380 effect; *F* _*1,68*_ = 13,69; *p* < 0.001***; O vs S1; Bonferonni’s multiple comparisons test: *Scrib*^*crc/+*^*-*CGP57380, *t*_*9*_ = 8.04, *p* < 0.0001***; Figure 5E). The same effects were observed on the preference for social novelty in treated conditions (no genotype x stimulus x CGP57380 effect; genotype x stimulus effect: *F* _*1, 62*_ = 6.18, *p* < 0.05*; S1 vs S2; Bonferonni’s multiple comparaisons test: *Scrib1*^*crc/+*^*-*CGP57380, *t*_*8*_ = 4.46, *p* < 0.0001***; Figure 5E). These data demonstrate that the impairments in social recognition observed in *Scrib*^*crc/+*^ mutant is in part due to an increase in ERK/Mnk1 pathway activity, consecutive to a reduction of Scrib levels in the hippocampus. Further, we demonstrate in our mouse model that social interest deficit can be rescued in adult animal by modulating two biological targets which are part of a major signaling pathway.

## Discussion

This study provides extensive evidence for the role of Scrib in the regulation of social behaviors, mediated by hippocampal functionality and -more specifically-by the ERK/Mnk signaling pathway. Three major findings were obtained from this study. First, we report that *Scrib*^*crc/+*^ mice have behavioral alterations that are specific to the social domain, in the absence of changes in other abilities. Second, these social deficits are associated with an increased c-Fos activity in the hippocampus and an over-activation of phosphorylated ERK. Third, the social abnormalities of *Scrib*^*crc/+*^ mice could be rescued by the administration of MEK/ERK/Mnk1 inhibitors. Our study identifies that the hyperactivation of ERK and Mnk and the dysregulation of the ERK/Mnk1 signaling pathways in the absence of a fully functional Scrib protein is the key molecular mechanism that triggers social behavior defects in the *Scrib*^*crc/+*^ mutant mice. Taken together, our findings suggest that Scribble controls ERK and Mnk1 signaling molecule levels of phosphorylation during social behavior.

### Scrib mutation induces behavioral alterations that are specifically social

*Scrib*^*crc/+*^ mice displayed lack of social interest and of preference for social novelty and social reward in the three-compartment test, as well as reduced social habituation. The qualitative patterns of social interaction, as assessed in the direct social interaction test, were unaltered, as well as ultrasonic communication. As we performed mainly quantitative measures of ultrasonic frequency and duration, it is still possible that subtle differences in communication may be revealed, for instance through qualitative spectrographic analyses as suggested by studies on mouse models of ASD (Scattoni et al., 2013). Nonetheless, our findings undoubtedly indicate that social recognition and detection of social novelty represent the major domains that are robustly affected by Scrib *crc* mutation. Based on these results, we have therefore focused on these social domains to identify the underlying neuromolecular mechanisms. The use of the three-compartment test further simplifies the procedures to be applied for the future and it enhances reproducibility across studies, by applying automatic measurements of compartment preference, thus avoiding the potential confounding impact of inter-observer variations of direct interaction analysis.

Importantly, the social deficits of *Scrib*^*crc/+*^ mouse mutant were not accompanied by other behavioral alterations. This allowed us to rule out the confounding role of olfactory, emotional and sensory motor alterations, but also to exclude a general novelty discrimination deficit, as supported by the lack of abnormalities in the Y maze and object recognition tests. Our data therefore suggest that the role of *SCRIB* in the etiopathology of ASD would be specific for social alterations, without other additional sensori-motor and emotional symptoms or cognitive deficits. This does not undermine the importance of the Scrib *crc* mouse line to investigate ASD and NDDs on the contrary. First, the social phenotype represents the most relevant core symptom of ASD-related pathologies and it is common to a variety of NDDs. Interestingly, other polarity genes have been linked with NDDs or more specifically ASD-like phenotypes (Sans et al., 2016). Notably, mutations in core PCP genes of the Dishevelled (*DVL*) or Prickle (*PK*) families are found in autistic patients (Dong et al., 2014; Gilman et al., 2011; Sowers et al., 2013), while the transgenic mouse model with mutations in two Disheveled genes, Dvl1 and Dvl3, display social behavioral abnormalities (Belinson et al., 2016; Gilman et al., 2011). Thus, our findings underscore the interest in studying the specific link between other polarity genes and social behaviors. Second, it is possible that Scrib may be linked to specific types of ASD or NDDs, for instance the high functioning or Asperger’s syndromes: this hypothesis could be supported by the higher spatial memory abilities previously described in the water maze task in Scrib mutant mice (Moreau et al., 2010), and would deserve further investigation.

### A central role of the hippocampus in Scrib *dependent* social dysfunctions

Using MRI analysis, we found no major structural alterations in the brain of our mutants except larger olfactory bulbs and a smaller hippocampus. Our data are in agreement with studies showing that the hippocampus plays a critical role in social memory and recognition (Kogan et al., 2000; Maaswinkel et al., 1996; Okuyama, 2018; Raam et al., 2017; Tzakis and Holahan, 2019). It is possible that social memories also rely on different subregions of the hippocampus, different networks and connections and/or different molecular pathways. More than one of them could be affected in the *circletail* mutant. It has recently been shown that a brain network composed of the hippocampus/mPFC/ACC/amygdala was required for the consolidation of social recognition memory (Tanimizu et al., 2017). In the *Scrib*^*crc/+*^ mice, both the hippocampus and amygdala showed an excessive c-Fos induction, a result that might highlight an abnormal regulation of neuronal activity resulting in deficient social memory to form properly. Moreover, MRI studies have highlighted reduced volume of amygdala or hippocampus, in children and adult ASD patients compared to typical subjects (Stanfield et al., 2008; Via et al., 2011) as well as in adult patients with spina bifida (Treble-Barna et al., 2015). It will be interesting to further explore the role of the hippocampus in these different forms of memory.

### The ERK/Mnk1 pathway mediates the impact of Scrib on social behavior: a novel therapeutic target?

Our data show that social exposure leads to an increased neuronal and MAPK signaling activity of the hippocampus in Scrib^*crc/+*^ mice compared to WT mice. Though various signaling pathway have been associated with ASD, this major signaling pathway is linked to neurodevelopmental disorders and more specifically ASD (Courchesne et al., 2019; Faridar et al., 2014; Vithayathil et al., 2018) and the ERK MAPK/ERK pathway is hyperactive in at least a subpopulation of autism spectrum disorder (Kalkman, 2012). Because Scrib levels are reduced in *Scrib*^*crc/+*^ mice, it is reasonable to think that this hyper-phosphorylation of ERK1/2 is the results of a release of inhibition of the MAPK cascade. This hypothesis is largely supported by studies is different tissue and cell types, demonstrating that full length Scrib inhibits ERK phosphorylation by interacting directly with ERK and inhibiting its subsequent translocation to the nucleus (Dow and Humbert, 2007; Nagasaka et al., 2013). Our luciferase assay demonstrated that MAPK activity reduction was dose-dependent, as more Scrib lead to more inhibition. We further found a biphasic regulation of the mutated *Scrib*^*crc*^ form of Scrib, suggesting a tight and dynamic regulation of MAPK activity depending on the levels of Scrib but also on the domains of the proteins available in neurons. Similar complex effect of a mutation is observed in a neuroligin-3 knock-in mouse, with a R451C-substitution likely acting as a gain-of-function mutation (Etherton et al., 2011). In this study, the authors observed that although only 10% of the neuroligin-3 protein remains in the knock-in mice, this remaining neuroligin-3 protein lead to an inhibition of synaptic transmission whereas the full neuroligin-3 KO exerts no such effect. Consistent with this, chromosomal duplications/deletions of 16p11.2, which includes the MAPK3 gene encoding for ERK1, or a microdeletion on chromosome 22, which contain the MAPK1 gene coding for ERK2 are commonly found associated with neurodevelopmental disorders and ASD (Campbell et al., 2008; Fernandez et al., 2010; Saitta et al., 2004).

We demonstrate in our mouse model that social interest deficit can be rescued in adult animal by modulating two biological targets which are part of a major signaling pathway. Mnk has emerged over the years as an important target in the ERK dependent signaling pathway that could be targeted to prevent ASD (Bramham et al., 2016). A recent study further suggests that MAPK activity might constitute a molecular signature of clinical severity in autism (Rosina et al., 2019). In that study, the authors suggest that the cascade ERK1-2/Mnk1/eIF4E may constitute a molecular signature predictive for early diagnosis of autism, notably severe ASD. Our results showed that inhibiting Mnk1 has similar effects than inhibiting directly ERK phosphorylation, both on sociability and on social novelty behavior, strongly suggesting it is a similar cascade that is deregulated in the *Scrib*^*crc/+*^ mice.

Alternatively, even if not mutually exclusive, Scrib could negatively regulate ERK activity indirectly, through the regulation of Rac1 activity. Many studies showed that Rac1 acts upstream of ERK1/2 through a p21-activated kinase (PAK) PAK/Raf/MEK signaling cascade, while *Scrib*^*crc/+*^ mice display enhanced Rac1 activity in the hippocampus (Moreau et al., 2010). Scrib could therefore be a negative regulator of Rac1 signaling in the hippocampus. Since Rac1 can activate ERK1/2, decreased levels of Scrib would also lead to increased activation of pERK (Kennedy et al., 2005). Consistent with this hypothesis, Tonegawa and collaborators were able to rescue symptoms of Fragile X syndrome – another major developmental disorder often associated to autism - in mice using a dominant negative PAK transgene (Hayashi et al., 2007). PAK also acts downstream of Scrib (Montcouquiol et al., 2006b; Sans et al., 2016), so it is reasonable to think that these proteins belong to the same molecular pathway, which appear as a good candidate for drugs that could improve social symptoms. Interestingly, PAK is also downstream of βPIX and SHANK proteins, which are main players in autistic behaviors with and without mental retardation (Park et al., 2003). Because of many similar phenotypes in *SHANK3* mouse models and in *Scrib*^*crc/+*^ mice, it would be of interest to evaluate whether Scrib and Shank3 act in concert or in parallel upstream of the ERK pathway to regulate social behavior.

### Conclusion

Altogether, this work identifies Scrib as an important regulator of social behavior in mice, through negative regulation of the ERK/Mnk signaling pathway, and that the *circletail* mutant mouse is a useful model on which the effects on social behavior of candidate therapeutical compounds could be tested. It is tempting to suggest that disruption of SCRIB-dependent signaling could underlie common genetic factors to NTD and ASD-like deficits, and may in some way predispose to development of these pathologies. In the future, it will be important to define the upstream and the other downstream effectors of Scrib implicated in the processing of the sensory inputs that play a pivotal role in social information integration.

## Materials and Methods

### Animals

Experiments were performed using heterozygous mice for the mutation (*Scrib*^*crc/+*^) and WT (*Scrib*^*+/+*^) littermates. All subjects were adult males of 10-11 weeks of age at the start of behavioral tests and housed in collective cages under standard laboratory conditions with a 12 h light/12 h dark cycle (light on: 07:00) with food and water supplied *ad libitum*. Juvenile male Swiss mice (21–28 days old, Janvier, France), or SV27 Female mice (10-12 weeks) were used as social stimulus for social interaction and communication analysis. Multiple cohorts of mice were used for the study, as described in the supplementary Materials and Methods section.

### Ethics Statement

Experiments were approved by the local Animal Health and Care Committee and performed in strict compliance with the EEC recommendations for the care and use of laboratory animals.

### Assessment of Social behaviours

The multiple social tests used assessed multiple characteristics of social behaviors in different social contexts. All data from the three-compartment test were analyzed with Ethovision (Version 13, Noldus). *Social interest and preference for social novelty in the three-compartment test*. The three-compartment test provided an evaluation of a social preference without mandatory direct social interaction; compared to direct social interaction tests, it allows to assess social behaviours in at context of a free choice, also avoiding inter-male aggression or sexual behaviours. The test was performed as previously described (Moreau et al., 2010). It consisted of three 10-min trials. During trial #1 (habituation), the tested mouse was allowed to explore the 3-chamber in which each end-chamber contained an empty small wire cage. In trial #2 (social interest), a stranger male mice (S1, social stimulus) was placed under a wire cage in one end-chambers while an object (O, non-social stimulus) was placed in the opposite end-chamber. For the trial #3 (social novelty), a second stranger male mice (S2) replaced the object. The tested mouse was free to choose between a caged novel stranger (S2, novel social stimulus) versus the same caged mouse in trial 2 (S1, familiar social stimulus).

#### Preference for social reward in the three-compartment test

The mouse was habituated to the 3-compartment apparatus for 10 min in the empty compartment. Then, we tested the social reward over 5 min. We placed a novel adult male stranger mouse (M) under a cage in one end-chamber and an adult female stranger mice (F) was placed in the opposite end-chamber. The tested mouse was free to choose between a both stimuli (F or M). Exploration was assessed by automatically measuring the time spent in each contact area containing the stimulus cage during 5 min.

#### Direct social interaction and communication

The quality of the behavioural patterns of social behaviours were assessed using the direct social interaction test with an adult female in in the home-cage. This experimental setting is also typically used to analyze ultrasonic vocalizations in adult male mice (Portfors, 2007). Tested mice were isolated 24 hours before the test. An unfamiliar stimulus mouse (adult NMRI female) was introduced into the testing cage and left for 5 min. Testing sessions were recorded and videos were analyzed with Observer XT (Noldus), taking only the tested mice into account. An observer (blind to the animal genotype) scored the time spent performing each of the following social behavior: sniffing the head and the snout of the partner, its anogenital region, or any other part of the body. During the test an ultrasonic microphone (CM16/CMPA, Avisoft, Berlin, Germany) was suspended 2 cm above the cage lid. Vocalizations were recorded and afterwards analyzed as previously described (Pietropaolo et al., 2011).

#### Social habituation

Social habituation was analyzed through repeated encounters with a juvenile male to elicit social interest but minimize the risk of aggression. The test was performed on two consecutive days. On day 1, an unfamiliar stimulus mouse (a juvenile Swiss male, 3-4 weeks old) was introduced in the home cage of the tested subject and left for 2 min. This procedure was repeated on day 2 with the same stimulus mouse. Social investigation of the stimulus mouse by the tested subject was scored by a trained observer who timed the duration of the investigation with a hand-held stopwatch. Behaviors that were scored as social investigation included direct contact, sniffings and close following (< 1cm).

### Assessment of non-social behaviors

Non-social tests for locomotion, emotionality, repetitive behaviors, olfactory discrimination/habituation, sensory-motor responsiveness and cognitive abilities including were evaluated as described in the supplementary section.

### Treatments

The MEK inhibitor α-[amino[(4-aminophenyl)thio]methylene]-2-(trifluoromethyl)benzeneacetonitrile (SL327) (Sigma-Aldrich) was dissolved in a vehicle solution of 2.5% dimethyl sulfoxide (DMSO)/ 2.5% Cremophor EL saline solution at 30 mg/kg (Satoh et al., 2011). The Mink1 inhibitor CGP57380 (Sigma-Aldrich) was dissolved in a vehicle solution of 2.5% dimethyl sulfoxide (DMSO) at 10mg/Kg (Lim et al., 2013). SL327 and CGP57380 were injected intraperitoneally (IP) at a volume of 3.3 ml/kg one hour before social interest test. Control mice received the same volume of vehicle.

### c-Fos immunoreactivity and laser microdissection following the social interest test

For each genotype, we analyzed two groups of mice: one group exposed to the social interest test (habituation and social interest = social group) and one group unexposed to the social interest test (habituation alone = control group). The social group went through the 2 trials of the three-compartment test described previously (trial 1 + trial 2). During the same period, the control group was exposed to the habituation test (trial1) during the same time in the same experimental room. At the end of the test, all mice were returned in their home cage and left undisturbed for 60 minutes (min).

#### c-Fos and Zif-268 immunoreactivity

Mice were anesthetized using a Phenobarbital i.p injection and transcardially perfused with physiological saline for 5 min followed by 4% paraformaldehyde diluted in 0.1 M phosphate buffer pH 7.4 (PFA4%) for 15 min. The brain was removed, postfixed for 24 h in 4% PFA and cut with a vibratome. Free-floating coronal sections (40 µm) were incubated in 0.3% Triton X-100, 0.3% normal goat serum and c-fos or Zif-268 primary antibody (Santa Cruz Biotechnology). Next, the sections were incubated with biotinylated goat anti-rabbit IgG (Vector Laboratories) followed by ABC EliteKit (Vector Laboratories). Immunoreactive cells were visualized using a diaminobenzidine (DAB, Vector Laboratories) colorimetric reaction. Images were acquired with a Leica microscope and the number of c-Fos and Zif-268 positive cells was quantified automatically using Metamorph. Brain areas were identified using the stereotaxic atlas of Paxinos and Franklin (1997) (Paxinos and Franklin, 2019).

#### Laser Capture Microdissection (LCM) analysis

Mice were anesthetized by brief inhalation of isoflurane (5% in air), sacrificed by decapitation and the brain was rapidly dissected, snap frozen, and stored at - 80°C. LCM was performed on coronal frozen sections as described (Maitre et al., 2011). The anterior hippocampal neuronal layers (B = -.94 à −2.7 mm) including the CA3, the CA1 and the DG were selectively captured on three different LCM caps. Following LCM, 150 µl of extraction buffer was added to the caps and stored at −80°C until protein isolation.

#### Western blot analysis

Preparation for protein extracts was performed as previously described (Moreau et al., 2010). The concentration of protein was measured by a BCA assay kit and western blots were done on 10% SDS-PAGE using standard methods (Sans et al., 2005). Antibodies used were anti-Scrib antibody (AbMM468 (Montcouquiol et al., 2006a); anti-p44/42 MAPK (#9102) and anti-Phospho-p44/42 MAPK (#9101) from Cell Signaling. Secondary antibodies used was anti-rabbit HRP-conjugated from GE Healthcare.

### Cell Culture, Transfection, and the SRE-Luciferase Reporter Assay

HEK293T cells were maintained in Dulbecco’s modified Eagle’s medium (GIBCO) supplemented with 10% fetal bovine serum (GIBCO) and were transfected using linear polyethylenimine (MW 25,000; Polysciences). SRE-Luciferase Reporter Assay was performed using the Dual-Luciferase assay system (Promega) as described previously (Ossipova et al., 2009). Each well in 12-well plates received 0.05 µg of pTKRenilla-Luc, 1 µg of pGL4.33[*luc2P*/SRE/Hygro] Vector DNA (Promega), and indicated amounts of human Scrib constructs (pCDNA3-eGFP-Scrib, pCDNA3-eGFP-Scrib^Crc^). Total DNA was adjusted to 2.05 µg by supplementing pCDNA3 DNA. One day after transfection, cells were serum starved for 2 hr then treated with 20% FBS containing medium for an additional 6 hr and harvested in 200µl of 1X passive lysis buffer for luciferase activity measurement. Results are expressed in terms of relative luciferase activity (ratio of firefly luciferase activity divided by Renilla luciferase activity), which were determined using a POLARstar Omega Multi-Mode Microplate Reader (BMG LABTECH).

### Magnetic resonance imaging (MRI)

Experiments were performed on a 4.7T Bruker system (Ettlingen, Germany) equipped with a 12-cm gradient system capable of 660 mT/m maximum strength. Measurements were performed with a birdcage resonator (25 mm diameter and 30 mm length) tuned to 200 MHz. Mice were anesthetized with isoflurane (1-1.5 % in air) and maintained at a constant respiration rate of 75 ± 15 respirations/min. The animals were placed in a lying position within the magnet with the head at the centre of the NMR coil. A 3D TrueFISP imaging sequence with alternating RF phase pulse method and sum of square reconstruction were used as already described (Miraux et al., 2008; Ribot et al., 2015). The following parameters were used: TE/TR: 2.5/5 ms; flip angle: 30°; bandwidth: 271 Hz/pixel; FOV: 20×20×16 mm; matrix: 128×128×102; spatial resolution: 156×156×156 mm^3^); number of averages: 4; 4 DeltaPhi values; total acquisition time: 17 min 04 s.

### 3D volume reconstitution and surface rendering

Consecutive z-series of coronal brain sections were stained with cresyl violet and scanned with Hamamatsu NANOZOOMER 2.0. All brain images were compiled into a stack file after being aligned using ImageJ software. 3D reconstruction and volume measurement were done with Imaris Scientific 3D/4D image processing and analysis software.

### Statistical analysis

All animals were assigned randomly to the different experimental conditions. Normality and homogeneity of variances from the samples were tested with Shapiro-Wilk normality test and Bartlett test respectively. If data were parametric, Student’s *t-test* (two comparisons), ANOVA 1-way (multiple comparisons), ANOVA 2-way or ANOVA 3-way were used. Otherwise non-parametric tests were used (Mann-Whitney or Kruskall-Wallis). When significant interaction effects of main factors were detected, post hoc analyses, recommended by GraphPad Prism 8 software (Bonferroni’s for ANOVA and Dunn’s for Kruskall-Wallis), were performed. Effects with p ≤ .05 was considered statistically significant.

## Funding and Disclosure

This research was supported by the INSERM, the University of Bordeaux, the CNRS, the Region Aquitaine and the ANR (ANR SAMENTA SynChAUTISM (ANR-13-SAMA-0012) to Y.C and NS). B.J.A.R was supported by a INSERM-Region Aquitaine PhD fellowship. The authors declare no conflict of interest.

## Acknowledgements

We thank M. Biguery for making the 3-compartment maze, L. Micheau for the help with behavioral tests, M.C. Donat, L. Lasvaux, R. Peyroutou and C. Medina for technical assistance and Dr. J.M. Revest for the gift of the ERK antibodies. We thank the animal and genotyping facilities of the Neurocentre Magendie for technical assistance, and notably M. Jaquet, H. Doat, V. Charbonnier, A.-L. Huot, F. Corailler, L. Dupuy, and D. Gonzales and co-workers. We thank Y. Rufin and the Biochemistry and Biophysics Platform of the Bordeaux Neurocampus at the Bordeaux University for the western-blot analysis. We thank C. Pujol and the team at the Bordeaux Imaging Center (BIC, a service unit of the CNRS-INSERM & Bordeaux Univ., member of France BioImaging, national infrastructure supported by the French National Research Agency (ANR-10-INSB-04, “Investments for the Future”) for the Metamorph macro and the technical help. All these facilities are funded by the LabEX BRAIN ANR-10-LABX-43. We finally, we thank Dr Claudia Racca for helpful comments and discussion.

## Author Contributions and information

MMM, SP, WEC, MM and NS conceptualized the paper. MMM, SP, JE, BJAR, SM and MaM participated in the execution and analysis of experiments. MMM, SP, JE, NS designed and interpreted the results. MMM, SP, MM and NS wrote the paper; All authors edited and approved the paper. Equal senior authorship: Mireille Montcouquiol and Nathalie Sans

## Notes

### Competing Interest Statement

The authors have declared no competing interest.

## References

Adolphs, R. (2009). The Social Brain: Neural Basis of Social Knowledge. Annu. Rev. Psychol. 60, 693–716.

Bagni, C., and Zukin, R.S. (2019). A Synaptic Perspective of Fragile X Syndrome and Autism Spectrum Disorders. Neuron 101, 1070–1088.

Belinson, H., Nakatani, J., Babineau, B., Birnbaum, R., Ellegood, J., Bershteyn, M., McEvilly, R., Long, J., Willert, K., Klein, O., et al. (2016). Prenatal β-catenin/Brn2/Tbr2 transcriptional cascade regulates adult social and stereotypic behaviors. Mol. Psychiatry 21, 1417–1433.

Bonello, T.T., and Peifer, M. (2019). Scribble: A master scaffold in polarity, adhesion, synaptogenesis, and proliferation. J. Cell Biol. 218, 742–756.

Bramham, C.R., Jensen, K.B., and Proud, C.G. (2016). Tuning Specific Translation in Cancer Metastasis and Synaptic Memory: Control at the MNK-eIF4E Axis. Trends Biochem. Sci. 41, 847–858.

Campbell, D.B., Li, C., Sutcliffe, J.S., Persico, A.M., and Levitt, P. (2008). Genetic evidence implicating multiple genes in the MET receptor tyrosine kinase pathway in autism spectrum disorder. Autism Res. Off. J. Int. Soc. Autism Res. 1, 159–168.

Cheng, N., Alshammari, F., Hughes, E., Khanbabaei, M., and Rho, J.M. (2017). Dendritic overgrowth and elevated ERK signaling during neonatal development in a mouse model of autism. PloS One 12, e0179409.

Courchesne, E., Pramparo, T., Gazestani, V.H., Lombardo, M.V., Pierce, K., and Lewis, N.E. (2019). The ASD Living Biology: from cell proliferation to clinical phenotype. Mol. Psychiatry 24, 88–107.

Dauber, A., Golzio, C., Guenot, C., Jodelka, F.M., Kibaek, M., Kjaergaard, S., Leheup, B., Martinet, D., Nowaczyk, M.J.M., Rosenfeld, J.A., et al. (2013). SCRIB and PUF60 are primary drivers of the multisystemic phenotypes of the 8q24.3 copy-number variant. Am. J. Hum. Genet. 93, 798–811.

Dawson, S., Glasson, E.J., Dixon, G., and Bower, C. (2009). Birth Defects in Children With Autism Spectrum Disorders: A Population-based, Nested Case-Control Study. Am. J. Epidemiol. 169, 1296–1303.

Diaz-Beltran, L., Esteban, F.J., and Wall, D.P. (2016). A common molecular signature in ASD gene expression: following Root 66 to autism. Transl. Psychiatry 6, e705.

Dong, S., Walker, M.F., Carriero, N.J., DiCola, M., Willsey, A.J., Ye, A.Y., Waqar, Z., Gonzalez, L.E., Overton, J.D., Frahm, S., et al. (2014). De novo insertions and deletions of predominantly paternal origin are associated with autism spectrum disorder. Cell Rep. 9, 16–23.

Dow, L.E., and Humbert, P.O. (2007). Polarity regulators and the control of epithelial architecture, cell migration, and tumorigenesis. Int. Rev. Cytol. 262, 253–302.

Dulawa, S.C., Grandy, D.K., Low, M.J., Paulus, M.P., and Geyer, M.A. (1999). Dopamine D4 receptor-knock-out mice exhibit reduced exploration of novel stimuli. J. Neurosci. Off. J. Soc. Neurosci. 19, 9550–9556.

Etherton, M., Földy, C., Sharma, M., Tabuchi, K., Liu, X., Shamloo, M., Malenka, R.C., and Südhof, T.C. (2011). Autism-linked neuroligin-3 R451C mutation differentially alters hippocampal and cortical synaptic function. Proc. Natl. Acad. Sci. U. S. A. 108, 13764–13769.

Farbman, A.I. (1994). The cellular basis of olfaction. Endeavour 18, 2–8.

Faridar, A., Jones-Davis, D., Rider, E., Li, J., Gobius, I., Morcom, L., Richards, L.J., Sen, S., and Sherr, E.H. (2014). Mapk/Erk activation in an animal model of social deficits shows a possible link to autism. Mol. Autism 5, 57.

Fernandez, B.A., Roberts, W., Chung, B., Weksberg, R., Meyn, S., Szatmari, P., Joseph-George, A.M., Mackay, S., Whitten, K., Noble, B., et al. (2010). Phenotypic spectrum associated with de novo and inherited deletions and duplications at 16p11.2 in individuals ascertained for diagnosis of autism spectrum disorder. J. Med. Genet. 47, 195–203.

Fernandez, S.M., Lewis, M.C., Pechenino, A.S., Harburger, L.L., Orr, P.T., Gresack, J.E., Schafe, G.E., and Frick, K.M. (2008). Estradiol-induced enhancement of object memory consolidation involves hippocampal extracellular signal-regulated kinase activation and membrane-bound estrogen receptors. J. Neurosci. Off. J. Soc. Neurosci. 28, 8660–8667.

Gilman, S.R., Iossifov, I., Levy, D., Ronemus, M., Wigler, M., and Vitkup, D. (2011). Rare de novo variants associated with autism implicate a large functional network of genes involved in formation and function of synapses. Neuron 70, 898–907.

Guillot, P.V., and Chapouthier, G. (1996). Olfaction, GABAergic neurotransmission in the olfactory bulb, and intermale aggression in mice: modulation by steroids. Behav. Genet. 26, 497–504.

Hayashi, M.L., Rao, B.S.S., Seo, J.-S., Choi, H.-S., Dolan, B.M., Choi, S.-Y., Chattarji, S., and Tonegawa, S. (2007). Inhibition of p21-activated kinase rescues symptoms of fragile X syndrome in mice. Proc. Natl. Acad. Sci. U. S. A. 104, 11489–11494.

Hilal, M.L., Moreau, M.M., Racca, C., Pinheiro, V.L., Piguel, N.H., Santoni, M.-J., Dos Santos Carvalho, S., Blanc, J.-M., Abada, Y.-S.K., Peyroutou, R., et al. (2017). Activity-Dependent Neuroplasticity Induced by an Enriched Environment Reverses Cognitive Deficits in Scribble Deficient Mouse. Cereb. Cortex N. Y. N 1991 27, 5635–5651.

Hu, J., Sathanoori, M., Kochmar, S., Azage, M., Mann, S., Madan-Khetarpal, S., Goldstein, A., and Surti, U. (2015). A novel maternally inherited 8q24.3 and a rare paternally inherited 14q23.3 CNVs in a family with neurodevelopmental disorders. Am. J. Med. Genet. A. 167A, 1921–1926.

Iossifov, I., Ronemus, M., Levy, D., Wang, Z., Hakker, I., Rosenbaum, J., Yamrom, B., Lee, Y.-H., Narzisi, G., Leotta, A., et al. (2012). De novo gene disruptions in children on the autistic spectrum. Neuron 74, 285–299.

Jarjour, A.A., Boyd, A., Dow, L.E., Holloway, R.K., Goebbels, S., Humbert, P.O., Williams, A., and ffrench-Constant, C. (2015). The polarity protein Scribble regulates myelination and remyelination in the central nervous system. PLoS Biol. 13, e1002107.

Kalkman, H.O. (2012). Potential opposite roles of the extracellular signal-regulated kinase (ERK) pathway in autism spectrum and bipolar disorders. Neurosci. Biobehav. Rev. 36, 2206–2213.

Kennedy, M.B., Beale, H.C., Carlisle, H.J., and Washburn, L.R. (2005). Integration of biochemical signalling in spines. Nat. Rev. Neurosci. 6, 423–434.

Kogan, J.H., Frankland, P.W., and Silva, A.J. (2000). Long-term memory underlying hippocampus-dependent social recognition in mice. Hippocampus 10, 47–56.

Lauritsen, M.B., Mors, O., Mortensen, P.B., and Ewald, H. (2002). Medical disorders among inpatients with autism in Denmark according to ICD-8: a nationwide register-based study. J. Autism Dev. Disord. 32, 115–119.

Lei, Y., Zhu, H., Duhon, C., Yang, W., Ross, M.E., Shaw, G.M., and Finnell, R.H. (2013). Mutations in planar cell polarity gene SCRIB are associated with spina bifida. PloS One 8, e69262.

Lim, S., Saw, T.Y., Zhang, M., Janes, M.R., Nacro, K., Hill, J., Lim, A.Q., Chang, C.-T., Fruman, D.A., Rizzieri, D.A., et al. (2013). Targeting of the MNK-eIF4E axis in blast crisis chronic myeloid leukemia inhibits leukemia stem cell function. Proc. Natl. Acad. Sci. U. S. A. 110, E2298–2307.

Lin, Y.-C., Frei, J.A., Kilander, M.B.C., Shen, W., and Blatt, G.J. (2016). A Subset of Autism-Associated Genes Regulate the Structural Stability of Neurons. Front. Cell. Neurosci. 10, 263.

Maaswinkel, H., Baars, A.-M., Gispen, W.-H., and Spruijt, B.M. (1996). Roles of the basolateral amygdala and hippocampus in social recognition in rats. Physiol. Behav. 60, 55–63.

Maitre, M., Roullot-Lacarrière, V., Piazza, P.V., and Revest, J.-M. (2011). Western blot detection of brain phosphoproteins after performing Laser Microdissection and Pressure Catapulting (LMPC). J. Neurosci. Methods 198, 204–212.

Miraux, S., Massot, P., Ribot, E.J., Franconi, J.-M., and Thiaudiere, E. (2008). 3D TrueFISP imaging of mouse brain at 4.7T and 9.4T. J. Magn. Reson. Imaging JMRI 28, 497–503.

Montcouquiol, M., Rachel, R.A., Lanford, P.J., Copeland, N.G., Jenkins, N.A., and Kelley, M.W. (2003). Identification of Vangl2 and Scrb1 as planar polarity genes in mammals. Nature 423, 173–177.

Montcouquiol, M., Sans, N., Huss, D., Kach, J., Dickman, J.D., Forge, A., Rachel, R.A., Copeland, N.G., Jenkins, N.A., Bogani, D., et al. (2006a). Asymmetric localization of Vangl2 and Fz3 indicate novel mechanisms for planar cell polarity in mammals. J. Neurosci. Off. J. Soc. Neurosci. 26, 5265–5275.

Montcouquiol, M., Crenshaw, E.B., and Kelley, M.W. (2006b). Noncanonical Wnt signaling and neural polarity. Annu. Rev. Neurosci. 29, 363–386.

Moreau, M.M., Piguel, N., Papouin, T., Koehl, M., Durand, C.M., Rubio, M.E., Loll, F., Richard, E.M., Mazzocco, C., Racca, C., et al. (2010). The planar polarity protein Scribble1 is essential for neuronal plasticity and brain function. J. Neurosci. Off. J. Soc. Neurosci. 30, 9738–9752.

Moy, S.S., Nadler, J.J., Perez, A., Barbaro, R.P., Johns, J.M., Magnuson, T.R., Piven, J., and Crawley, J.N. (2004). Sociability and preference for social novelty in five inbred strains: an approach to assess autistic-like behavior in mice. Genes Brain Behav. 3, 287–302.

Murdoch, J.N., Henderson, D.J., Doudney, K., Gaston-Massuet, C., Phillips, H.M., Paternotte, C., Arkell, R., Stanier, P., and Copp, A.J. (2003). Disruption of scribble (Scrb1) causes severe neural tube defects in the circletail mouse. Hum. Mol. Genet. 12, 87–98.

Nagasaka, K., Massimi, P., Pim, D., Subbaiah, V.K., Kranjec, C., Nakagawa, S., Yano, T., Taketani, Y., and Banks, L. (2010). The mechanism and implications of hScrib regulation of ERK. Small GTPases 1, 108–112.

Nagasaka, K., Seiki, T., Yamashita, A., Massimi, P., Subbaiah, V.K., Thomas, M., Kranjec, C., Kawana, K., Nakagawa, S., Yano, T., et al. (2013). A novel interaction between hScrib and PP1γ downregulates ERK signaling and suppresses oncogene-induced cell transformation. PloS One 8, e53752.

Neale, B.M., Kou, Y., Liu, L., Ma’ayan, A., Samocha, K.E., Sabo, A., Lin, C.-F., Stevens, C., Wang, L.-S., Makarov, V., et al. (2012). Patterns and rates of exonic de novo mutations in autism spectrum disorders. Nature 485, 242–245.

Okuyama, T. (2018). Social memory engram in the hippocampus. Neurosci. Res. 129, 17–23.

Ossipova, O., Ezan, J., and Sokol, S.Y. (2009). PAR-1 phosphorylates Mind bomb to promote vertebrate neurogenesis. Dev. Cell 17, 222–233.

Park, E., Na, M., Choi, J., Kim, S., Lee, J.-R., Yoon, J., Park, D., Sheng, M., and Kim, E. (2003). The Shank family of postsynaptic density proteins interacts with and promotes synaptic accumulation of the beta PIX guanine nucleotide exchange factor for Rac1 and Cdc42. J. Biol. Chem. 278, 19220–19229.

Paxinos, G., and Franklin, K.B.J. (2019). Paxinos and Franklin’s the Mouse Brain in Stereotaxic Coordinates (Academic Press).

Pietropaolo, S., Guilleminot, A., Martin, B., D’Amato, F.R., and Crusio, W.E. (2011). Genetic-background modulation of core and variable autistic-like symptoms in Fmr1 knock-out mice. PloS One 6, e17073.

Portfors, C.V. (2007). Types and functions of ultrasonic vocalizations in laboratory rats and mice. J. Am. Assoc. Lab. Anim. Sci. JAALAS 46, 28–34.

Raam, T., McAvoy, K.M., Besnard, A., Veenema, A.H., and Sahay, A. (2017). Hippocampal oxytocin receptors are necessary for discrimination of social stimuli. Nat. Commun. 8, 2001.

Ribot, E.J., Wecker, D., Trotier, A.J., Dallaudière, B., Lefrançois, W., Thiaudière, E., Franconi, J.-M., and Miraux, S. (2015). Water Selective Imaging and bSSFP Banding Artifact Correction in Humans and Small Animals at 3T and 7T, Respectively. PloS One 10, e0139249.

Robinson, A., Escuin, S., Doudney, K., Vekemans, M., Stevenson, R.E., Greene, N.D.E., Copp, A.J., and Stanier, P. (2012). Mutations in the planar cell polarity genes CELSR1 and SCRIB are associated with the severe neural tube defect craniorachischisis. Hum. Mutat. 33, 440–447.

Rosina, E., Battan, B., Siracusano, M., Di Criscio, L., Hollis, F., Pacini, L., Curatolo, P., and Bagni, C. (2019). Disruption of mTOR and MAPK pathways correlates with severity in idiopathic autism. Transl. Psychiatry 9, 1–10.

Saitta, S.C., Harris, S.E., McDonald-McGinn, D.M., Emanuel, B.S., Tonnesen, M.K., Zackai, E.H., Seitz, S.C., and Driscoll, D.A. (2004). Independent de novo 22q11.2 deletions in first cousins with DiGeorge/velocardiofacial syndrome. Am. J. Med. Genet. A. 124A, 313–317.

Sans, N., Wang, P.Y., Du, Q., Petralia, R.S., Wang, Y.-X., Nakka, S., Blumer, J.B., Macara, I.G., and Wenthold R.J. (2005). mPins modulates PSD-95 and SAP102 trafficking and influences NMDA receptor surface expression. Nat. Cell Biol. 7, 1179–1190.

Sans, N., Ezan, J., Moreau, M.M., and Montcouquiol, M. (2016). Chapter 13 - Planar Cell Polarity Gene Mutations in Autism Spectrum Disorder, Intellectual Disabilities, and Related Deletion/Duplication Syndromes. In Neuronal and Synaptic Dysfunction in Autism Spectrum Disorder and Intellectual Disability, C. Sala, and C. Verpelli, eds. (San Diego: Academic Press), pp. 189–219.

Satoh, Y., Endo, S., Nakata, T., Kobayashi, Y., Yamada, K., Ikeda, T., Takeuchi, A., Hiramoto, T., Watanabe, Y., and Kazama, T. (2011). ERK2 contributes to the control of social behaviors in mice. J. Neurosci. Off. J. Soc. Neurosci. 31, 11953–11967.

Scattoni, M.L., Martire, A., Cartocci, G., Ferrante, A., and Ricceri, L. (2013). Reduced social interaction, behavioural flexibility and BDNF signalling in the BTBR T+tf/J strain, a mouse model of autism. Behav. Brain Res. 251, 35–40.

Sowers, L.P., Loo, L., Wu, Y., Campbell, E., Ulrich, J.D., Wu, S., Paemka, L., Wassink, T., Meyer, K., Bing, X., et al. (2013). Disruption of the non-canonical Wnt gene PRICKLE2 leads to autism-like behaviors with evidence for hippocampal synaptic dysfunction. Mol. Psychiatry 18, 1077–1089.

Stanfield, A.C., McIntosh, A.M., Spencer, M.D., Philip, R., Gaur, S., and Lawrie, S.M. (2008). Towards a neuroanatomy of autism: a systematic review and meta-analysis of structural magnetic resonance imaging studies. Eur. Psychiatry J. Assoc. Eur. Psychiatr. 23, 289–299.

Tanimizu, T., Kenney, J.W., Okano, E., Kadoma, K., Frankland, P.W., and Kida, S. (2017). Functional Connectivity of Multiple Brain Regions Required for the Consolidation of Social Recognition Memory. J. Neurosci. Off. J. Soc. Neurosci. 37, 4103–4116.

Timonen-Soivio, L., Sourander, A., Malm, H., Hinkka-Yli-Salomäki, S., Gissler, M., Brown, A., and Vanhala, R. (2015). The Association Between Autism Spectrum Disorders and Congenital Anomalies by Organ Systems in a Finnish National Birth Cohort. J. Autism Dev. Disord. 45, 3195–3203.

Treble-Barna, A., Juranek, J., Stuebing, K.K., Cirino, P.T., Dennis, M., and Fletcher, J.M. (2015). Prospective and episodic memory in relation to hippocampal volume in adults with spina bifida myelomeningocele. Neuropsychology 29, 92–101.

Tzakis, N., and Holahan, M.R. (2019). Social Memory and the Role of the Hippocampal CA2 Region. Front. Behav. Neurosci. 13, 233.

Via, E., Radua, J., Cardoner, N., Happé, F., and Mataix-Cols, D. (2011). Meta-analysis of gray matter abnormalities in autism spectrum disorder: should Asperger disorder be subsumed under a broader umbrella of autistic spectrum disorder? Arch. Gen. Psychiatry 68, 409–418.

Vithayathil, J., Pucilowska, J., and Landreth, G.E. (2018). ERK/MAPK signaling and autism spectrum disorders. Prog. Brain Res. 241, 63–112.

Wang, M., Marco, P. de, Capra, V., and Kibar, Z. (2019). Update on the Role of the Non-Canonical Wnt/Planar Cell Polarity Pathway in Neural Tube Defects. Cells 8.

