## Supplementary Information for "Scribble controls social behaviors through the regulation of the ERK/Mnk1 pathway"

#### Supplementary materials and methods

***Animals.***

The mutant line has been maintained by random breeding between heterozygous *Scrib1*$^{crc/+}$ and wild-type *Scrib1*$^{+/+}$ mice. The genetic background of the transgenic animals line is 50% NMRI, 25% BALB/C, and 25% C57BL/6 (Murdoch et al., 2001). Several cohorts of animals and multiple behavioural tests were used. Whenever possible, naïve animals were employed for behavioural testing; when the same cohort was used for multiple tests, the most stressful essays were administered last, to minimize between-test interference.

One group (n = 8 WT and n = 16 *Scrib1*$^{crc/+}$) was tested in open field and in light dark test. Second group (n = 13 WT and n = 12 *Scrib1*$^{crc/+}$) was tested in plus maze, Y-maze, pre-pulse inhibition and social communication. Three naives group were used for social interest and preference for social novelty in the three-compartment test and olfactory tests (n = 9 WT and n = 10 *Scrib1*$^{crc/+}$), the preference for social reward (n = 7 WT and n = 11 *Scrib1*$^{crc/+}$) and the social habituation (n = 11 WT and n = 15 *Scrib1*$^{crc/+}$). Differences groups (n = 8-10 WT and n = 7-10 *Scrib1*$^{crc/+}$) was used for the ERK and Mnk1inhibitor experiments. One last group was used for self-grooming and marble test (n = 9 WT and n = 10 *Scrib1*$^{crc/+}$). All tests were performed during the light phase of the light/dark cycle. All experimental apparatuses were cleaned with Ethanol 70% or Phagospray DM between subjects to remove odour residuals. For others experiments (n = 3-4 WT and n = 5-6 *Scrib1*$^{crc/+}$) was used for c-Fos analysis; (n = 3 WT and n = 3 *Scrib1*$^{crc/+}$) was used for western blot, IRM and 3D analysis.

***Non-social tests for locomotion and emotionality.***

*Open field activity and non-social neophobia.* Open field activity was assessed during 30 min in brightly lit enclosures (40x40cm) equipped with infrared photocells to measure horizontal movements. Then, a novel object (a cup, 7.5 cm in height and 6.5 cm in diameter) was placed upside-down into the center of each open field and mouse behavior was tested for an additional 30 min. The percentage of entries and time spent in the center (15 cm in diameter), as well as the overall motor activity were quantified automatically by a software (Videotrack, Lyon, France).

*Elevated plus-maze.* The plus maze was made of grey Plexiglas and was elevated approximately 60 cm from the floor. It had two open and two enclosed arms radiating outward from a central open square. Mice were placed in the center of the maze and allowed free access to all arms for 5 min. The maze was illuminated by halogen light at 100 lux in the central open square. After each assessment, the equipment was cleansed with Phagospray DM to remove residual odors. Tracking images from a camera above the center of the apparatus were analyzed with Ethovision (Version 13, Noldus Technology, Wageningen, The Netherlands). The percentage time spent in the open arms was computed.

*Dark-light emergence test.* The emergence test (Dulawa et al., 1999) was conducted in a open field (MEASURES) containing a polypropylene cylinder (10cm-deep and 6.5 cm in diameter) located length wise along the wall, with the open located at a distance of 10 cm from the corner of the open field. Each subject was placed in the cylinder and left undisturbed in the apparatus for 15 mins. The latency to leave the cylinder, defined by the exit with all four paws into the open field, was measured as well as the number of entries and the time spent in the cylinder. Overall motor activity was quantified as the total distance performed during the test. The equipment was cleansed with Phagospray DM between animals to remove residual odors.

***Non-social tests for repetitive behaviors.***

*Self-grooming test*. Mice were scored for spontaneous grooming behaviors. WT or *Scrib1*$^{crc/+}$ mouse was placed individually in a new standard mouse cage with fresh bedding (46 cm length × 23.5 cm wide × 20 cm high, illuminated at 50 lux). After a 10-min habituation period in the test cage, each mouse was scored with a stopwatch for 10 min for cumulative time spent grooming all body regions.

*Marble burying test.* This test was considerate as an indicative measure of repetitive behavior in a novel environment related to digging behavior. The test consists in placing 20 glass marbles, spaced in five rows of four on approximately 5-cm layer of sawdust bedding, in a plastic cage. Mice were individually placed in the cage and after 30 min of test, the number of marbles buried with sawdust is counted. Marbles were considered buried if they were at least one half covered with bedding.

***Non-social tests for olfactory discrimination/habituation and sensory-motor responsiveness.***

*Buried food test without food deprivation.* On the day of the test, the mice were placed in a new cage (46 x 23.5 x 20 cm) containing 3 cm of bedding and allowed to explore for 10min. The animal was removed and a piece of cheese was buried under 2 cm of bedding. The time required to find the food was measured by an observer.

*Buried food test following food deprivation.* Mice were habituated to the flavor of a specific food (Kellogg's chocopops cereal) for 3 days before testing to avoid food neophobia on the day of testing. Twenty hours before the test, all food was removed from the home cage. On the day of the test, the mice were placed in a new cage (46 x 23.5 x 20 cm) containing 3 cm of bedding and allowed to explore for 10min. The animals were removed and a piece of cereal was buried under 2cm of bedding. The mice were returned in the cage and given 15 mins to locate the buried food. The time required to find the food was measured by an observer.

*Olfactory habituation and dishabituation.* For the test of olfactory habituation/dishabituation (adapted from Yang and Crawley 2009 (Yang and Crawley, 2009)), different odors (two non-social odors and two social odors) were presented in consecutive trials of 1 min each. First, the mouse was placed in the testing cage for habituation with a cotton applicator placed through a hole of the grid cage. After 30 min, a 10-µl drop of orange extract (Sanoflore, France; $10^{-6}$ dilution) was placed on the cotton-tip part of the applicator in the home cage for 1 min. The presentation of this odor stimulus was repeated three times with 10-min intervals. In a fourth trial 10 min later, we placed a new applicator containing 10-µl drop of lavandin extract (Sanoflore, France; $10^{-6}$ dilution) in the subject's cage for 1 min. Ten min later a piece of white paper containing male or female urine (a mixture of urine from 10 unfamiliar adult mice) were placed in the subject's cage sequentially in four consecutive trials of 1 min each. We considered olfactory investigation as the time sniffing the applicator.

*Prepulse inhibition of the acoustic startle reflex.* The apparatus (SR-LAB, San Diego Instruments, San Diego, CA, USA) and procedures were previously described in detail (Pietropaolo and Crusio, 2009; Pietropaolo et al., 2008a, 2008b; Yee et al., 2005) . Briefly, animals were acclimated to the apparatus for 5 min. The first six trials consisted of six pulse-alone trials, two for each pulse intensity (100, 110, or 120 dBA), presented in a pseudorandom order. Subsequently, ten blocks of trials were presented. Each block consisted of three pulse-alone trials, one for each pulse intensity, three prepulse-alone trials (+6, +12, or +18 dB units above the background of 65 dBA), nine possible combinations of prepulse-plus-pulse trials (3 levels of pulse × 3 levels of prepulse), and one no-stimulus trial (i.e., background alone). These 16 trials were presented in a pseudorandom order within each block, with a variable intertrial interval of a mean duration of 15 sec. The session was concluded with a final block of six consecutive pulse-alone trials as in the first block. Reactivity scores obtained on the first and the last blocks of six consecutive pulse-alone trials were separately analyzed to measure startle habituation. The data obtained in the remaining trials were categorized into three main different subsets. First, startle reactivity was assessed by the reactivity scores obtained in the intermediate pulse-alone trials. Second, reactivity on prepulse-plus-pulse trials relative to middle pulse-alone trials was used to estimate prepulse inhibition. PPI was analyzed converting the reactivity data into percent scores (%PPI = 100 × (pulse-alone − prepulse-plus-pulse)/pulse-alone) calculated for each subject for each of the nine possible prepulse-plus-pulse combinations. Third, to measure prepulse-elicited reactivity we included data from prepulse-alone and no-stimulus trials.

***Non-social tests for cognitive abilities.***

*Recognition of non-social novelty.* Each subject was placed in the center of an open field identical to the one previously described and allowed to explore during 10 min (Habituation phase). After habituation, two identical objects (orange china cup, 7.5 cm in height and 6.5 cm in diameter) were placed in two opposite corners of the open field and the experimental animal was left in the apparatus for 5 min (Sample phase). At the end of this session, the mouse was returned to his home cage. One hour after, one of the two objects was replaced by a novel once (grey plastic rectangle object: 4 × 8 cm) and the animal was given a 5-min exploration session (TEST PHASE). The time spent exploring each object (sniffing or/staying within 1cm distance) was measured during both sample and test phases by an observer.

*Spontaneous alteration in the Y maze.* Spontaneous alternation was assessed in a grey, plastic Y-maze, placed on a table 80 cm high and located in the middle of a room containing a variety of extramaze cues. The three arms of the Y-maze were similar in appearance and spaced at 120° from each other. Each arm was 42 cm long and 8 cm wide. The entire maze was enclosed by a wall 15 cm high and .5 cm thick. Mice were introduced at the end of one of the arms and allowed to explore the maze for 5 min. Allocation of the start arm was counterbalanced within experimental groups. An entry into one of the arms was scored by an observer unaware of the genotype of the animals when all four paws of the animal were placed inside an arm. Spontaneous alternation, expressed as a percentage, refers to that proportion of arm choices differing from the previous two (Hughes, 2004; King and Arendash, 2002). Thus, if an animal made the following sequence of arm choices: A, B, C, B, A, B, C, A, the total number of alternation opportunities would be six (total entries minus two) and the percentage alternation would be 67% (four out of six).

### Supplementary Tables

**■ Table S1, *Scrib1$^{crc/+}$* mice have normal basic motor functions, exploratory behavior, and anxiety**

| | WT | *Scrib1$^{crc/+}$* | *t*-test |
|---|---|---|---|
| Physical characteristics | | | |
| Body weight (gr) | 29.10 ± 0.55 | 28.50 ± 0.76 | $t_{30}$ = 0.61, n.s |
| Locomotor activity | | | |
| Open field : Total distance (cm/30min) | 9401.6 ± 434.4 | 10519.6 ± 6 | $t_{22}$ = 1.19, n.s |
| Plus maze : Total distance (cm/5min) | 807.2 ± 61.9 | 810.0 ± 50.3 | $t_{23}$ = 0.03, n.s |
| Repetitive behavior | | | |
| Spontaneous alternation Y maze (%) | 69.40 ± 2.55 | 70.60 ± 1.83 | $t_{23}$ = 0.37, n.s |
| Self-grooming (sec/10 min) | 63.87 ± 15.13 | 51.70 ± 10.03 | $t_{23}$ = 0.68, n.s |
| Rearing (sec/10 min) | 64.49 ± 8.676 | 64.34 ± 9.74 | $t_{23}$ = 0.011, n.s |
| Marble buried (number/30min) | 13.56 ± 1.88 | 14.70 ± 1.726 | $t_{23}$ = 0.44, n.s |
| Neophobia | | | |
| Time around the object (sec/30min) | 527.80 ± 113.3 | 429.80 ± 67.4 | $t_{22}$ = 0.78, n.s |
| Distance around the object (cm/30min) | 2529.5 ± 354.4 | 2058.5 ± 259.9 | $t_{22}$ = 1.05, n.s |
| Number of entries around the object | 67.25 ± 8.85 | 66.00 ± 5.57 | $t_{22}$ = 0.12, n.s |
| Anxiety | | | |
| Open field : Time in center (sec/30min) | 146.8 ± 11.5 | 148.8 ± 18.4 | $t_{22}$ = 0.07, n.s |
| Open field : Distance in center (cm/30min) | 1638.1 ± 152.1 | 1809.9 ± 227.2 | $t_{22}$ = 0.50, n.s |
| Plus maze : % Time in open arm (5min) | 15.69 ± 3.57 | 13.72 ± 2.70 | $t_{23}$ = 0.43, n.s |
| Plus maze : Number of entries (5min) | 15.77 ± 1.74 | 18.08 ± 1.81 | $t_{23}$ = 0.91, n.s |
| Light/Dark : latency to emerge (15min) | 112.13 ± 29.73 | 112.19 ± 56.01 | $t_{22}$ = 0.12, n.s |
| Light/Dark : Time inside (15min) | 342.25 ± 64.28 | 346.56 ± 54.37 | $t_{22}$ = 0.04, n.s |
| Light/Dark : Number of entries (15min) | 14.37 ± 1.73 | 16.62 ± 1.87 | $t_{22}$ = 0.76, n.s |

All data are presented as Mean ± SEM. No significant effect of genotype was observed (all n.s.)

**■ Table S2, *Scrib1*$^{crc/+}$ mice have normal pre pulse inhibition**

Arbitrary units for startle; %PPI for the rest:

| | | lnp100 | lnp110 | lnp120 | perc_PPI_100 | perc_PPI_110 | perc_PPI_120 |
|---|---|---|---|---|---|---|---|
| **WT** | mean | 2.43 | 3.42 | 4.54 | 13.41 | 33.24 | 26.95 |
| | SEM | 0.07 | 0.10 | 0.12 | 5.61 | 6.04 | 5.90 |
| ***Scrib1*$^{crc/+}$** | mean | 2.50 | 3.76 | 4.88 | 8.99 | 41.61 | 29.91 |
| | SEM | 0.12 | 0.16 | 0.15 | 7.49 | 4.82 | 5.70 |

**■ Table S3, Measure of change in Zif268 immunoreactive cells in response to social test (cells/mm² x10)**

| | **WT** | ***Scrib1*$^{crc/+}$** | ***p*** |
|---|---|---|---|
| **Unexposed** | | | |
| **DG** | 5.91 ± 1.45 | 4.97 ± 1.33 | *n.s* |
| **CA1** | 15,36 ± 3.77 | 17.59 ± 59 | *n.s* |
| **CA3** | 2.38 ± 1.10 | 3.65 ± 2.09 | *n.s* |
| **Social test** | | | |
| **DG** | 24.05 ± 2.62 | 39,54 ± 6.17 | $p < 0.05$ |
| **CA1** | 36.01 ± 3.48 | 36.74 ± 3.31 | *n.s* |
| **CA3** | 12.85 ± 1.25 | 22,65 ± 3.83 | $p < 0.05$ |

All data are presented as Mean ±SEM. * $p \leq 0.05$. No significant effect of genotype was observed (n.s.)

### Supplementary references

Dulawa, S.C., Grandy, D.K., Low, M.J., Paulus, M.P., and Geyer, M.A. (1999). Dopamine D4 receptor-knock-out mice exhibit reduced exploration of novel stimuli. J. Neurosci. Off. J. Soc. Neurosci. *19*, 9550–9556.

Hughes, R.N. (2004). The value of spontaneous alternation behavior (SAB) as a test of retention in pharmacological investigations of memory. Neurosci. Biobehav. Rev. *28*, 497–505.

King, D.L., and Arendash, G.W. (2002). Behavioral characterization of the Tg2576 transgenic model of Alzheimer's disease through 19 months. Physiol. Behav. *75*, 627–642.

Murdoch, J.N., Rachel, R.A., Shah, S., Beermann, F., Stanier, P., Mason, C.A., and Copp, A.J. (2001). Circletail, a new mouse mutant with severe neural tube defects: chromosomal localization and interaction with the loop-tail mutation. Genomics *78*, 55–63.

Pietropaolo, S., and Crusio, W.E. (2009). Strain-dependent changes in acoustic startle response and its plasticity across adolescence in mice. Behav. Genet. *39*, 623–631.

Pietropaolo, S., Feldon, J., and Yee, B.K. (2008a). Nonphysical contact between cagemates alleviates the social isolation syndrome in C57BL/6 male mice. Behav. Neurosci. *122*, 505–515.

Pietropaolo, S., Singer, P., Feldon, J., and Yee, B.K. (2008b). The postweaning social isolation in C57BL/6 mice: preferential vulnerability in the male sex. Psychopharmacology (Berl.) *197*, 613–628.

Yang, M., and Crawley, J.N. (2009). Simple behavioral assessment of mouse olfaction. Curr. Protoc. Neurosci. *Chapter 8*, Unit 8.24.

Yee, B.K., Chang, T., Pietropaolo, S., and Feldon, J. (2005). The expression of prepulse inhibition of the acoustic startle reflex as a function of three pulse stimulus intensities, three prepulse stimulus intensities, and three levels of startle responsiveness in C57BL6/J mice. Behav. Brain Res. *163*, 265–276.

### Supplementary figures

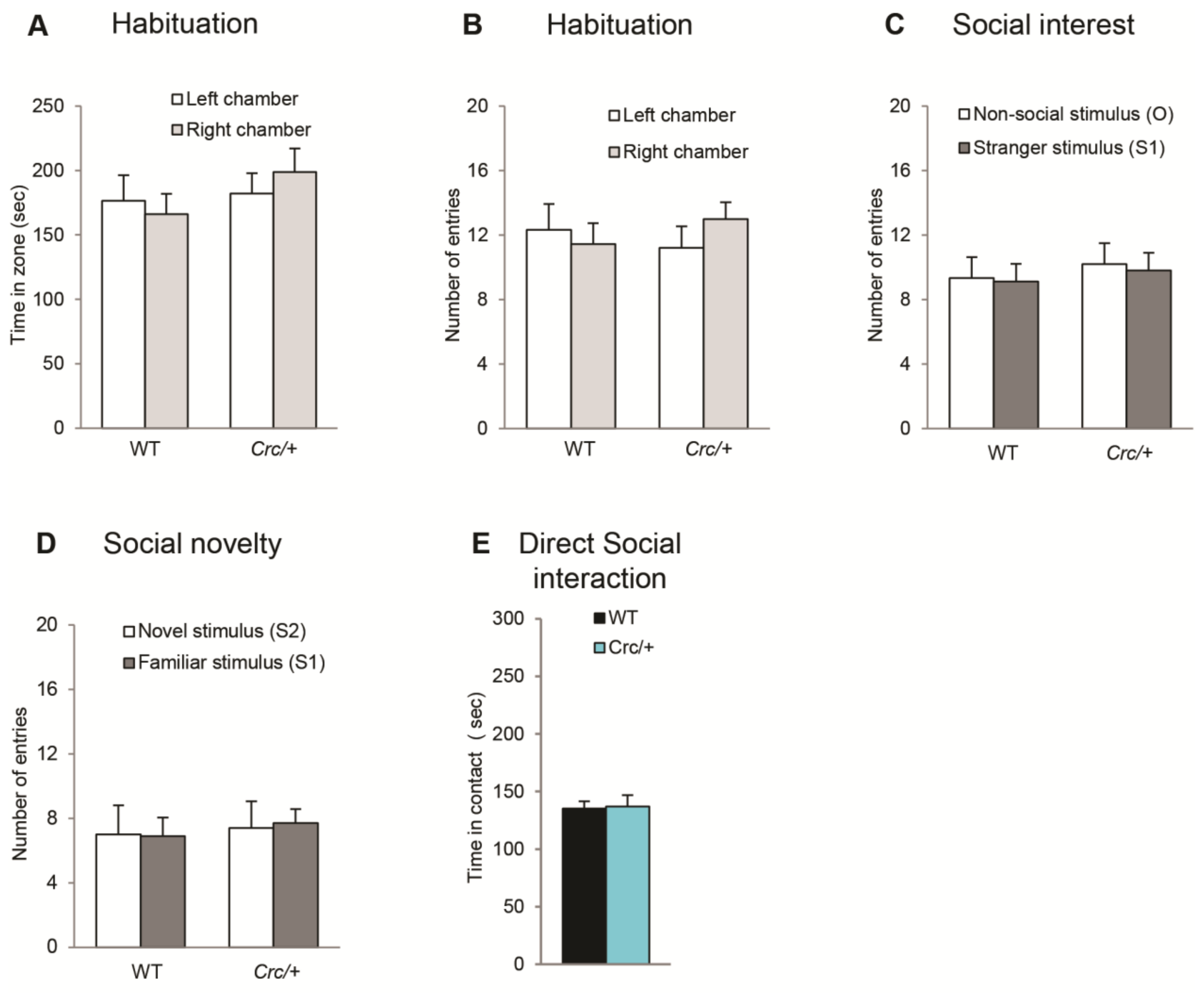


**Figure S1 The social behavior deficit of the *Scrib1*$^{crc/+}$ mice is not due to an activity deficit.** WT and *Scrib1*$^{crc/+}$ mice **a** spend the same time and **b** perform the same number of entries in the different chamber during the habituation test. WT and *Scrib1*$^{crc/+}$ mice perform the same number of entries in social stimulus chamber (S1) versus empty chamber (non-social stimulus, O) during **c** the social interest and **d** the Social novelty test. **e** WT and *Scrib1*$^{crc/+}$ mice perform the same number of contact direct social interaction test.

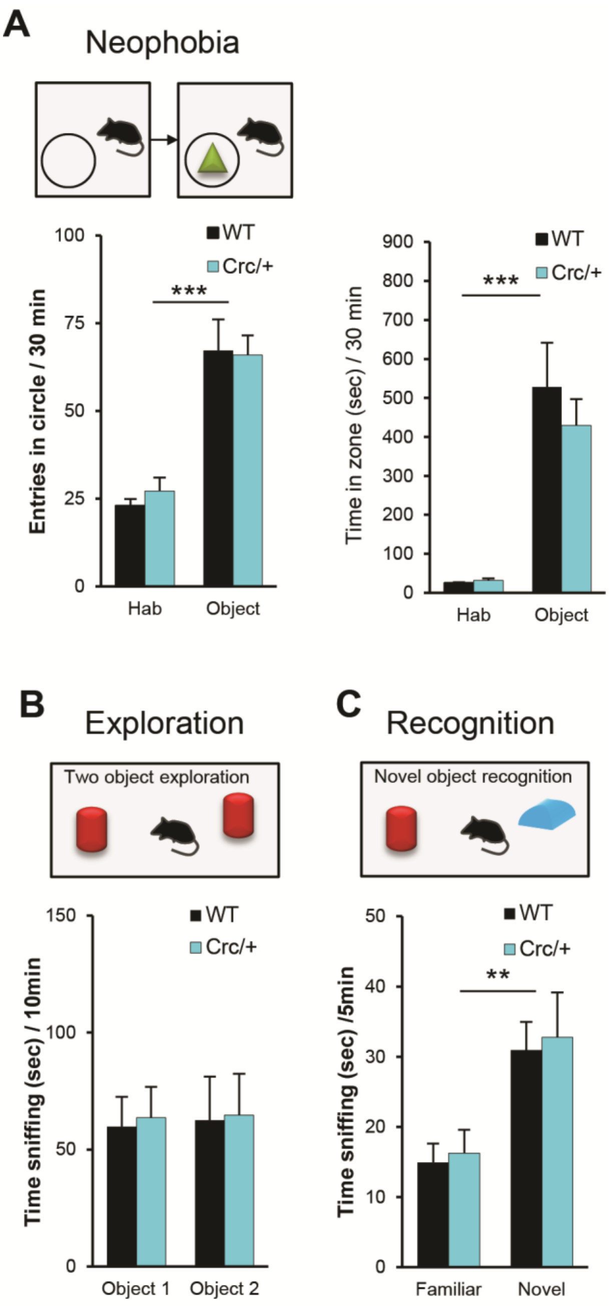


**Figure S2 The social behavior deficit of the *Scrib1*$^{crc/+}$ mice is not due to abnormal general novelty recognition. a** Experimental design protocol of the open field and neophobia test. Number of entries in the area which contain the object during the simple open field exploration (habituation) and the neophobia test (Object). WT and *Scrib1*$^{crc/+}$ mice explored equally the object. **b, c** Experimental design protocol of the two objects exploration and novel object recognition. **b** During training for object recognition, two objects were presented to a mouse during 10 min. WT and *Scrib1*$^{crc/+}$ mice spend the same time to explore the objects. **c** One hour after, one of the previous objects was changed by a novel object. Recognition memory measured the time spend sniffing the novel object versus the familiar object in the first 5 min of testing. Here two groups of mice presented no difference in recognition memory. All data are presented ±SEM. The link hook is used for significant object effect **$p \leq 0.01$ and ***$p \leq 0.001$.

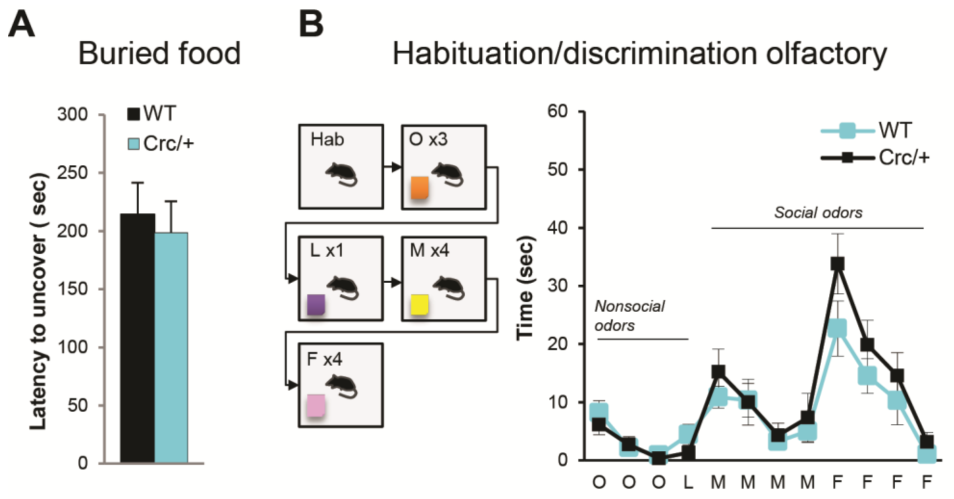


**Figure S3 The social behavior deficit of the *Scrib1*$^{crc/+}$ mice is not due to abnormal olfactory problems. a** Buried food test. The time spent to find the piece of cheese is the same for both mice groups, there is no difference between the genotype. **b** Experimental design protocol of habituation/dishabituation olfactory test. Habituation / dishabituation olfactory test was used to assess sense of smell. Non-social odor was presented following by social odors and mice were assessed for dishabituation on first presentation of novel odor and habituation on the third/four presentation of the odor. All data are presented ±SEM.

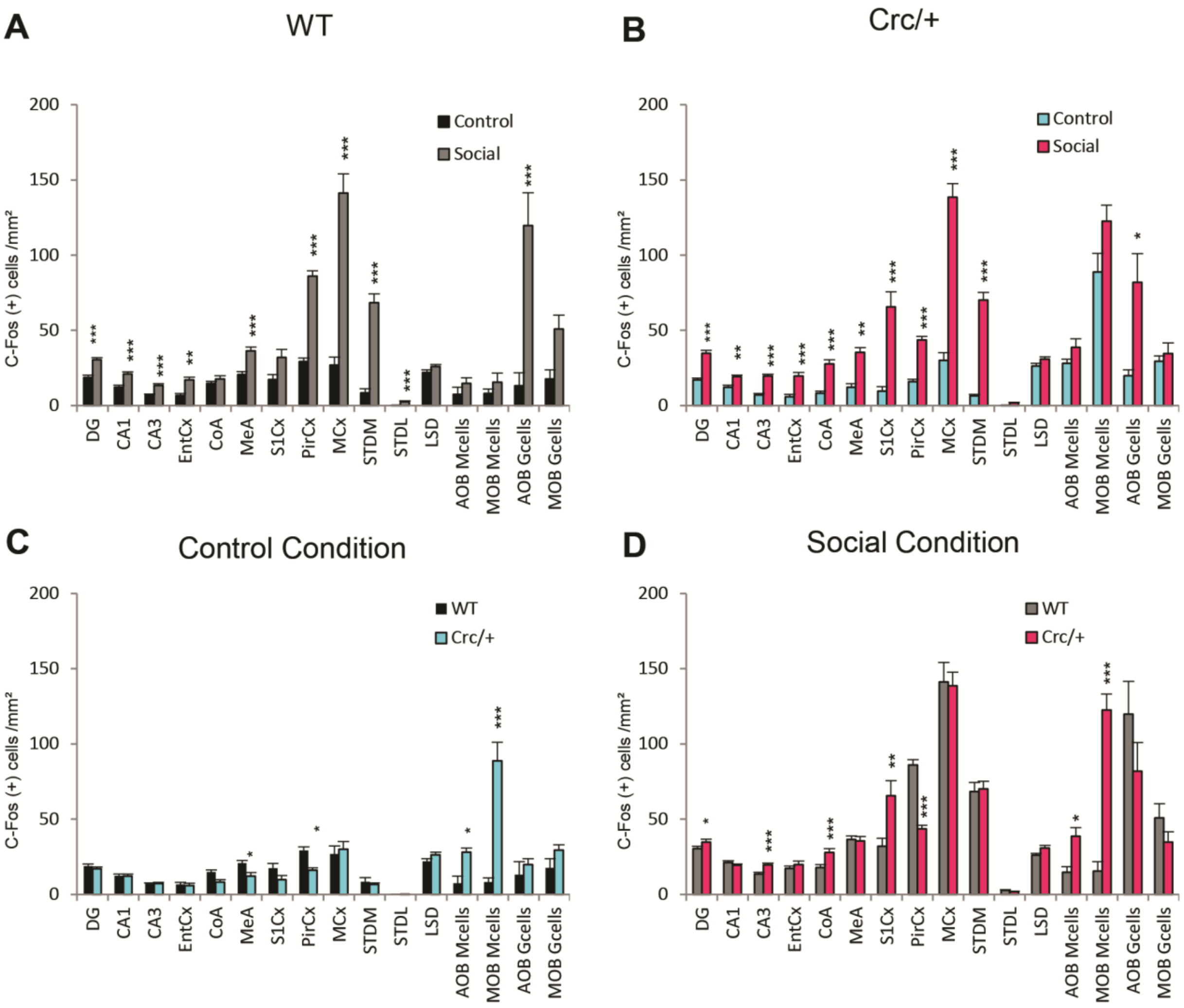


**Figure S4 Measure of change in c-Fos immunoreactive cells.** In response to social test **a** in the WT and **b** *Scrib1*$^{crc/+}$ mice. **c,d** Measure of change in c-Fos immunoractive cells in response to Scrib1 mutation in **c** the control condition and **d** within the social interest test. *$p \leq 0.05$; **$p \leq 0.01$ and ***$p \leq 0.001$.
